# Cognitive processing of a common stimulus synchronizes brains, hearts, and eyes

**DOI:** 10.1101/2021.09.16.460722

**Authors:** Jens Madsen, Lucas C. Parra

## Abstract

Neural, physiological and behavioral signals synchronize between human subjects in a variety of settings. Multiple hypotheses have been proposed to explain this interpersonal synchrony, but there is no clarity under which conditions it arises, for which signals, or whether there is a common underlying mechanism. We hypothesized that similar cognitive processing of a shared stimulus is the source of synchrony between subjects, measured here as inter-subject correlation. To test this we presented informative videos to participants in an attentive and distracted condition and subsequently measured information recall. Inter-subject correlation was observed for electro-encephalography, gaze position, pupil size and heart rate, but not respiration and head movements. The strength of correlation was co-modulated in the different signals, changed with attentional state, and predicted subsequent recall of information presented in the videos. There was robust within-subject coupling between brain, heart and eyes, but not respiration or head movements. The results suggest that inter-subject correlation is the result of similar cognitive processing and thus emerges only for those signals that exhibit a robust brain-body connection. While physiological and behavioral fluctuations may be driven by multiple features of the stimulus, correlation with other individuals is co-modulated by the level of attentional engagement with the stimulus.

## Introduction

It is well established that shared experiences can synchronize physiological signals between individuals. This has been observed for autonomic signals ^1^ including galvanic skin response, heart rate, body temperature, respiration, as well as other signals such gaze position ^2^ and pupil size ^3^. Synchronization of physiological signals between individuals has often been attributed to physical or social interaction ^4–7^. However, the simultaneous experience is not a prerequisite for physiological synchronization, as a number of studies have shown that physiological signals can be correlated between subjects even when they engage individually with dynamic natural stimuli ^3,8–11^.

Stimulus-induced correlation between individuals has been studied extensively with neural measures including fMRI ^12^, EEG ^13^, MEG ^14^ and fNIRS ^15^ in particular during presentation of film ^2,12^ and auditory narratives ^16,17^. These studies show that subjects process narrative stimuli similarly, and that correlation of brain activity is predictive of memory of the narrative ^17,18^. The cognitive state of viewers has been shown to influence the inter-subject correlation of neural activity. For instance, subjects that are attentive ^19^ and more engaged have higher high inter-subject correlation ^20^. In total, these studies indicate that perceptual and cognitive processing of the narrative are similar across subjects and depend on cognitive state.

In the context of physiological or behavioral signals, inter-subject correlation is often referred to as “interpersonal synchronization”, implying that two or more people are co-present in a given context. A variety of mechanisms have been proposed to cause inter-subject correlation such as social interactions ^1^, physical interactions ^4,7,21^, shared emotions ^9,11^, and it has been argued that the strength of synchrony is modulated by empathy ^22,23^, arousal ^4,24^, attention ^25^, and more ^1^. The diversity of factors parallels the diversity of factors known to affect physiological signals. For instance, heart rate fluctuations are often discussed in the context of emotions, and pupil size in the context of arousal, although both are affected by a variety of other cognitive factors ^26–32^. We hypothesize, instead, that the cognitive processing of a shared stimulus is sufficient to induce inter-subject correlation, thus providing a more parsimonious explanation for the variety of phenomena previously observed. We use the term “cognitive processing” in its general sense of creation and manipulation of mental representations, which includes stimulus perception ^33^. The hypothesis predicts that inter-subject correlation is co-modulated in different signals, and importantly, that it will emerge only for physiological signals that exhibit robust coupling to the brain. If, instead, physiological synchrony is driven by a variety of factors, we would not expect a co-modulation of ISC nor do we expect signals with robust brain-body coupling to necessarily synchronize between subjects.

To test these opposing predictions we collected physiological, neural and behavioral signals while participants watched informative videos. Data was collected individually for each participant to rule out effects of direct social interactions. Additionally, videos were selected to be engaging but not to evoke strong emotions or arousal as in previous studies on heart rate synchronization ^6,8,9^. These controls can falsify alternative hypotheses that require social or physical interactions ^1,4,7,21^ or strong emotions or empathy ^8,9,11,22,23^. Our hypothesis also predicts that inter-subject correlation should be modulated by attention and predictive of memory of the content in the video. To test for this we used a secondary mental task that distracts attention from the stimulus ^19^ and subsequently probed for recall memory ^34^ We measured inter-subject correlation (ISC) for each individual ^12,13^, and resolved it on the group level also across time and frequency.

As predicted, we find significant inter-subject correlation between individuals only for those signal modalities that exhibit a robust brain-body connection, namely, gaze position, pupil size, heart rate and saccade rate. We did not find significant correlation for respiration or head velocity, which indeed did not exhibit a robust coupling with brain activity. Consistent with our hypothesis, the strength of ISC co-varied across signal modalities, was modulated by attention and was predictive of recall. These results suggest that inter-subject correlation is modulated in unison by the level of attentional engagement with the stimulus.

## Results

To establish the strength of inter-subject correlation in different modalities we presented instructional videos while simultaneously recording neural, behavioral and physiological signals. In the first experiment with 92 subjects (Experiment 1) we recorded the electroencephalogram (EEG), heart rate (HR), gaze position (gaze), pupil size (pupil) and respiration. We chose 3 instructional videos related to physics, biology, and computer science, each 3-5 min long, with a total duration of 10 min. Signals were recorded at different sampling frequency for each modality but aligned in time across all subjects and modalities (Fig. 1A). For each signal modality we compute the Pearson’s correlation of these time courses between all pairs of subjects (Fig. 1B). For gaze position the correlation is computed separately for horizontal and vertical position and then averaged. For the 64 channels of EEG, we first extract several components of the raw evoked potentials that maximally correlate between subjects ^35^, compute pairwise correlation between subjects for each component, and then take the sum of the correlation values. In each signal modality, intersubject correlation (ISC) is defined for each subject as the average Pearson’s correlation between that subject and all others (Fig 1C).

**Figure 1.**
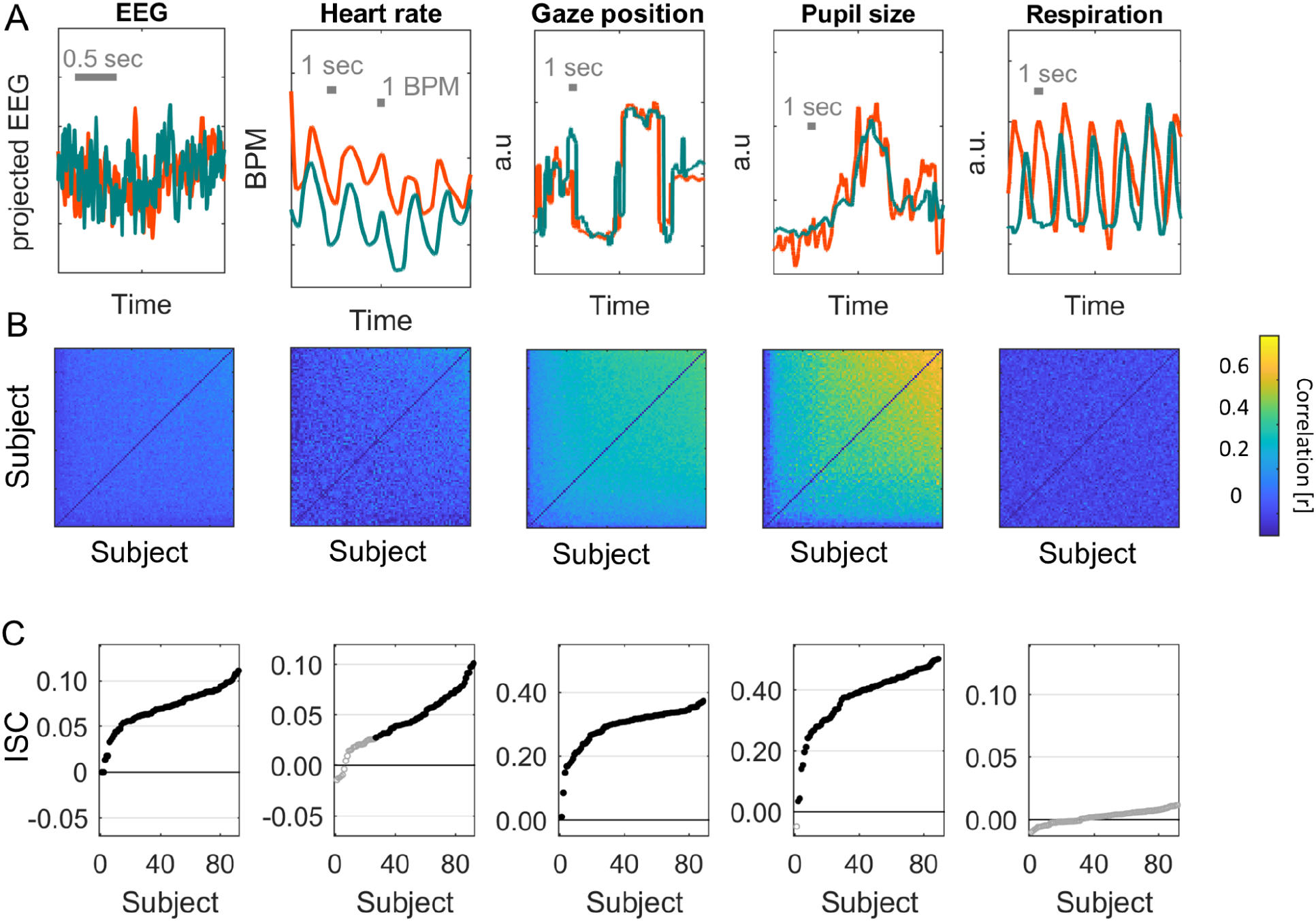
Inter-subject correlation of neural, physiological and behavioral signals during passive video watching. **A)** Signals for each of the modalities simultaneously recorded during Experiment 1. The two subjects shown (green and orange) have the highest ISC values measured for each modality. The EEG signal is the first component extracted from the 64 channel EEG using correlated component analysis. The gaze position signal is the horizontal gaze position. **B)** Pearson correlation matrix between all pairs of the 92 subjects for each of the modalities. Subjects are sorted by increasing average correlation values. The correlation matrix for gaze position is the average correlation of gaze position in the horizontal and vertical direction. The correlation matrix for EEG is the sum of the correlation values obtained for 9 components extracted with correlated component analysis. **C)** ISC values are the average of pairwise correlations for each subject, i.e. the mean over column of the correlation matrix in B, excluding the diagonal, and averaged over the 3 videos presented (10 min total duration). Subjects are ordered by their ISC values (same as in B). Filled points indicate statistically significant ISC values and non-filled points indicate they are not statistically significant. Statistical significance is determined using circular shuffle statistics (10,000 shuffles and corrected for multiple comparisons with FDR of 0.01). Circular shuffle means that the signal of each subject is randomly shifted in time, thus removing any intersubject relation.

The first observation is that ISC is statistically significant in most subjects and modalities (black points in Fig. 1C), but also quite variable across individuals. Significant correlation was detected in all modalities except respiration. Comparing ISC across modalities we see that the most robust correlation is found for gaze position (ISC_gaze_ is in the range of 0.01-0.37, mean M=0.29, standard deviation SD=0.06). It is significantly larger from chance values for all 92 subjects tested. Robust correlation is also found for pupil size (ISC_pupil_: 0.06-0.51, M=0.39, SD=0.09, 92/92 significant) and EEG (ISC_EEG_: 0.00–0.11, M=0.07, SD=0.02, 92/92 significant). We find that heart rate also correlates between subjects, but to a lesser extent with several subjects not exhibiting significant correlation (ISC_HR_: −0.01-0.10, M=0.04, SD=0.03, 61/92 significant). For respiration we were not able to detect significant correlation between any participant and the group (ISC_resp_: −0.01–0.01, M=0.00, SD=0.00).

### Inter-subject correlation co-varies across different modalities

Given the strong variation of ISC across subjects, we wanted to determine if it co-varies in different signal modalities, i.e. if ISC is high in one modality, is it also high in other modalities? To this end we compare the ISC of different modalities across subjects (Fig. 2). Subjects with high inter-subject correlation in EEG, heart rate, gaze position and pupil size also showed high correlation in the other signal modalities. Correlation of ISC across subjects between these four modalities is significant in all cases (r=0.46–0.72, p<0.01, N=92). However, we do not find any significant relation between the ISC of these four modalities and respiration (r=-0.07–0.06, p>0.1, N=92). The observation that the level of ISC of gaze position, pupil size and EEG co-varies across subjects is perhaps expected as they may all be driven by the visual dynamic of the video. What is more surprising is that high ISC in these modalities also coincided with high ISC of heart rate fluctuations, which thus far has been mostly attributed to emotional aspects of a stimulus ^8^ and would not be expected to be driven by visual dynamics of the video.

**Figure 2.**
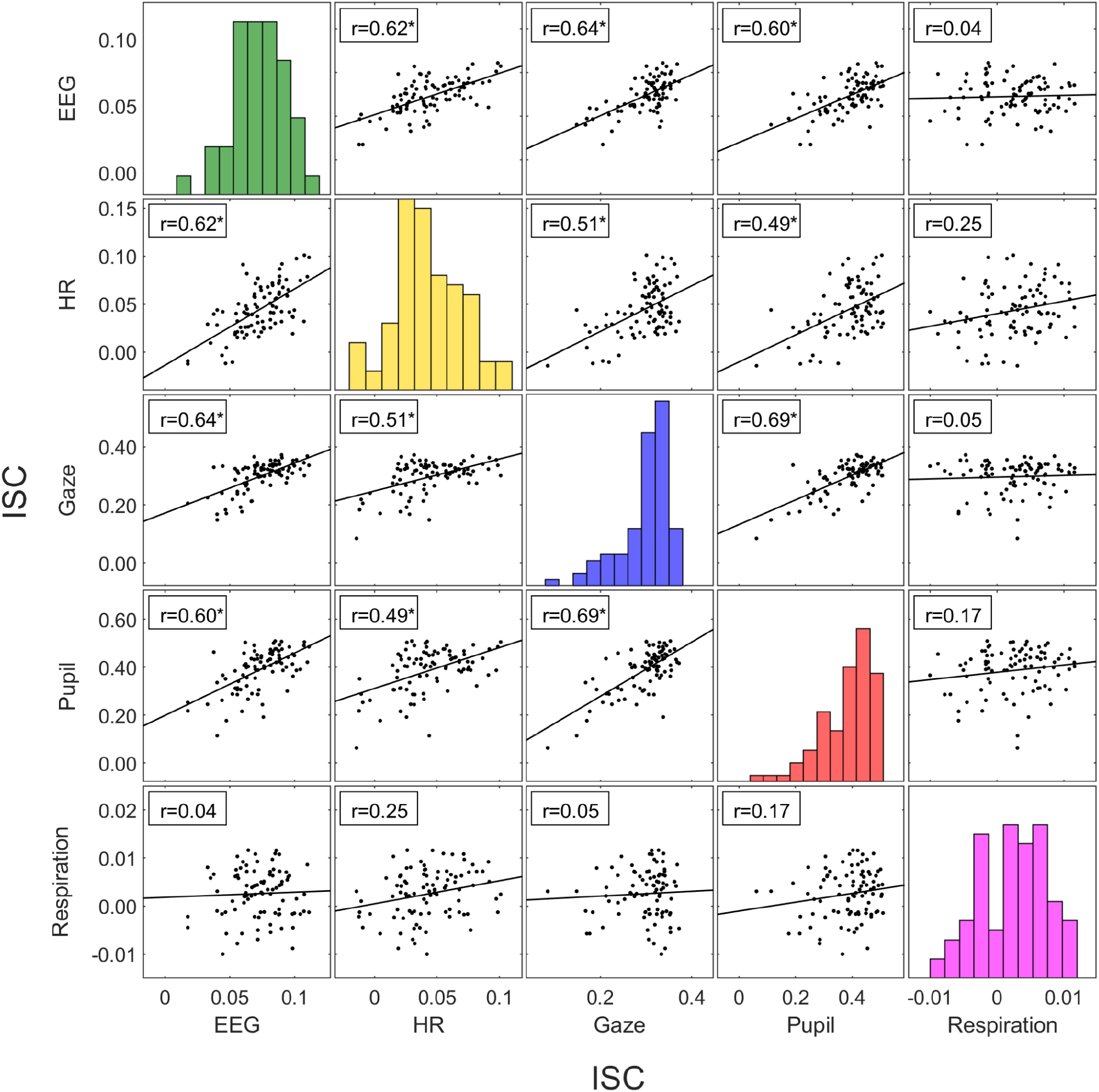
Intersubject correlation is co-modulated in different signal modalities. Inter-subject correlation computed here for multiple signals recorded on 92 subjects while they watched instructional videos (same as in Fig. 1). Each point represents ISC between one individual and the group. The diagonal of the plot matrix is the histogram of each modality. Significant Pearson’s correlation of ISC between each modality is indicated by a * (p<0.01, uncorrected). Lines indicate the linear least-squares prediction of vertical from horizontal axis (while points are the same in upper and lower triangles after flipping axes, prediction lines are not).

### Time scale of correlated signal fluctuations and co-modulation across time

ISC captures whether subjects move their eyes in unison, whether their heart rate increases or decreases together, whether their pupils dilate or contract or whether they inhale and exhale simultaneously. Given that ISC is co-modulated between modalities, we wanted to know if these correlated fluctuations were due to similar entrainment with the stimulus. To investigate this, we resolved ISC by frequency band, i.e. the signals are band-pass filtered prior to computing ISC resulting in a coherence spectrum. These coherence spectra differ significantly between modalities (Fig. 3A). Therefore, co-modulation is not likely to result from entrainment to a specific frequency band, and instead may be driven by more complex properties of the stimulus. While coherent fluctuations differ between modalities, they generally are slower than 10Hz and are strong in the frequency band around 0.1Hz. ISC might therefore be reliably measured on a time scale of 10s. We additionally analyzed the power spectra, which quantifies the magnitude of fluctuations in different frequency bands (Fig. 3B). They differ significantly from the coherence spectrum. Therefore coherence is frequency-specific and not just a result of the underlying dynamic. Given this diversity, it is possible that each signal modality is modulated by something different in the stimulus across time. We therefore asked whether ISC, from one time interval to the next, change together in different modalities. Indeed, time-resolved ISC of brain signals computed on 10s intervals correlates significantly with time-resolved ISC of gaze position, pupil size and heart rate (with correlations in the range of r=0.1-0.4), but not with respiration (Fig. 3C). This correlation of ISC between modalities over time is weaker than what we found across subjects.This suggests that on a short time scale (less than 15 min), ISC of different modalities may be driven by a diversity of factors in these video stimuli. Nevertheless, both across subjects and across time, we find that ISC of different signal modalities are co-modulated.

**Figure 3:**
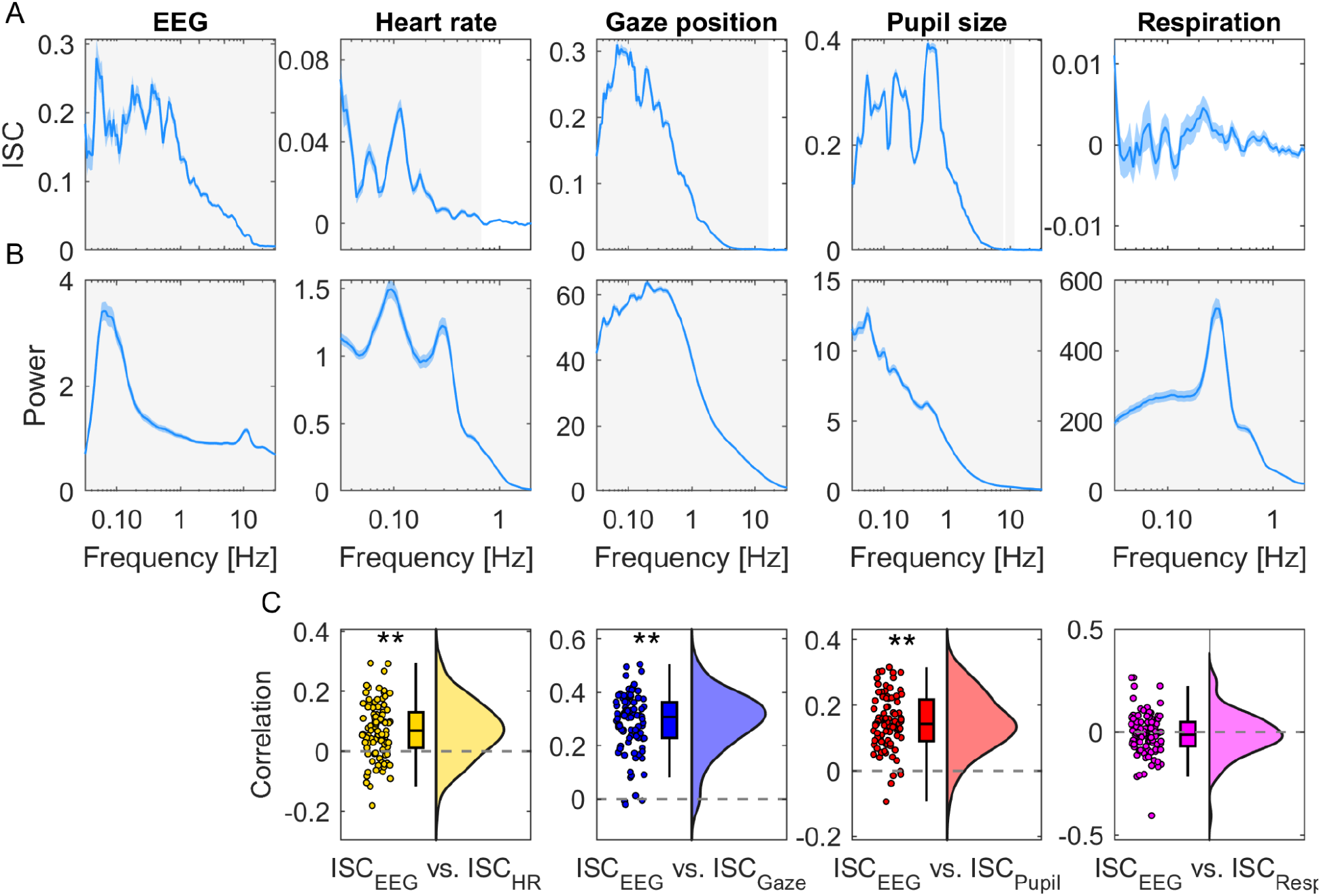
Time scales of signal fluctuations. **A)** Frequency-resolved ISC: Inter-subject coherence spectrum computed by first band-pass filtering signals at different frequencies and then computing ISC averaged over videos and subjects. Band-pass filtering used center frequency on a logarithmic scale and a bandwidth of 0.2 octaves. Blue shading indicates SEM across N=92 subjects. Significance of ISC values above zero are established in each band using t-test, corrected for multiple comparisons using one-dimensional cluster statistics (light grey area indicates p<0.01, cluster corrected). **B)** Magnitude of fluctuations captured by the power spectrum, i.e. the power of the band-pass filtered signals. **C)** Correlation between time-resolved ISC of EEG and time-resolved ISC of different modalities computed separately for each subject. Time-resolved ISC is computed in 10 second time windows with 50% overlap. Each dot is a subject. ** indicate p<0.01 for a t-test for non-zero mean ISC, uncorrected). The density is computed using kernel density estimation using a gaussian kernel. The box plot shows the median and 25th and 75th percentile. Data aggregated over ~15 min of video of Experiment 1.

### Inter-subject correlation is predictive of an individual’s memory performance

We hypothesized that inter-subject correlation is the result of cognitive processing, and thus we expected that the level of ISC will be predictive of memory performance. We therefore tested memory of the material in the video after presentation with a set of multiple choice questions (these data were available for 43 of the 92 subjects). Questions tested memory of the information presented in the video, which covered topics related to science, technology and math (STEM). Questions probed for recognition and comprehension such as “What are stars made of?” or “How can Single Nuclear Polymorphism be useful?”. First, to quantify the common factor that is driving the co-modulation of ISC across subjects, we used principal component analysis (after z-scoring ISC values for each modality). The first principal component captures 52% of the population variance in ISC and the second component only 22% (Fig. 4A). The modalities captured by this first component are EEG, HR, gaze position and pupil size (Fig. 4B), consistent with the co-variation of ISC observed in Fig. 2. Whereas the second principal component loaded mostly on respiration (Fig. 4B). As predicted, we find a strong correlation of memory performance with the first principal component of ISC (Fig. 4C) (r(39)=0.61, p=2.7· 10^-5^, gaze position data was missing in 2 of the 43 subjects). In contrast, there is no significant correlation with the second component and test taking performance (r(39)=-0.14, p=0.38).

**Figure 4.**
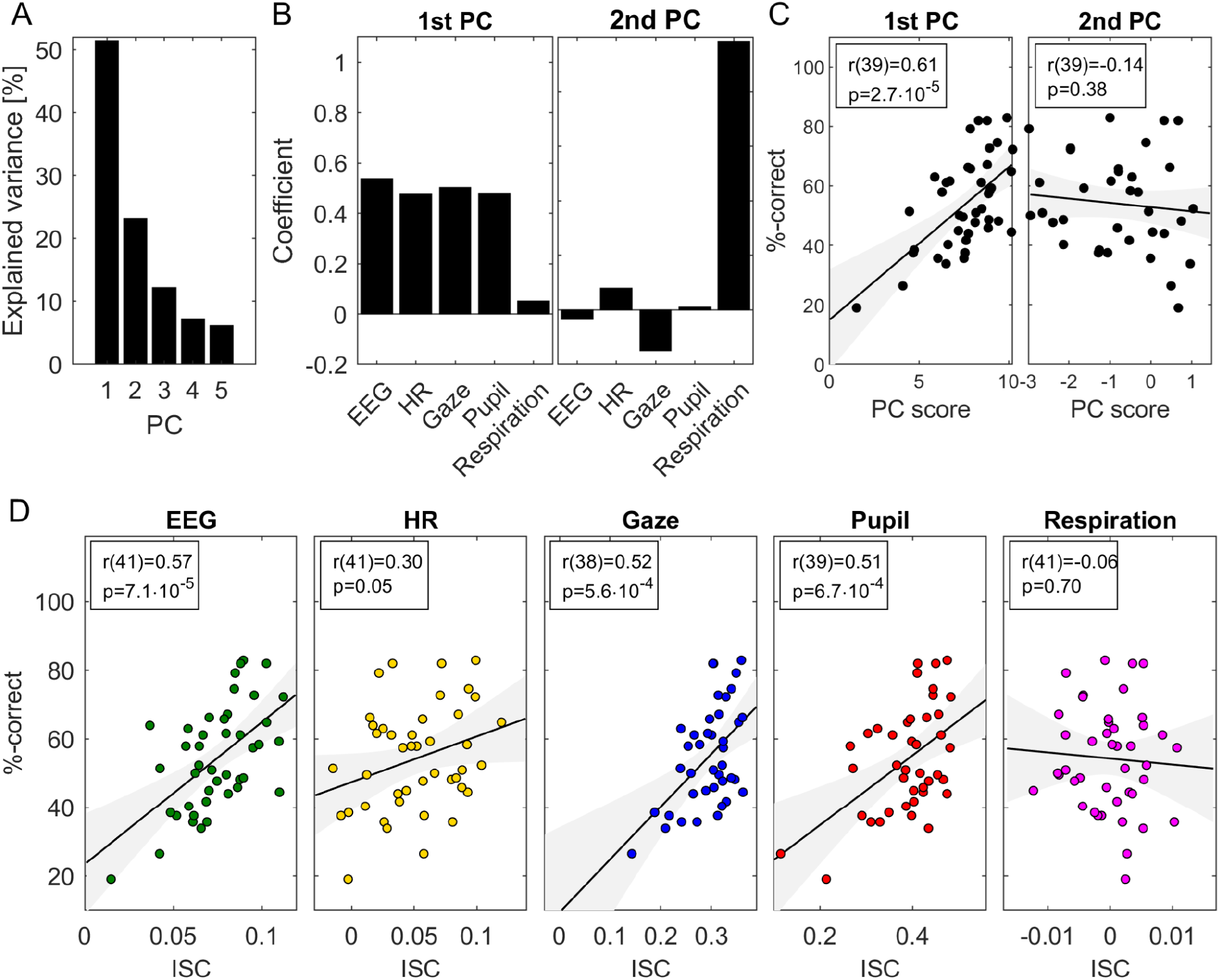
Common factor modulating ISC predicts memory recall: **A)** Variance explained by the principal components computed across all ISC modalities. **B)** Loadings on the first two principal components, i.e. how much of each ISC modality is captured by each of the principal components. **C)** The ISC projected onto the two first principal coefficients (PC Score) and the test taking performance of students (%-correct). **D)** ISC of each modality and the test taking performance of students. Panels A and B include all data from Fig. 3 (Experiment 1) Memory performance for panels C and D was available only for a subset of 43 subjects. P-values are uncorrected.

This result is confirmed when we look at each modality independently (Fig. 4D). ISC of EEG is predictive of students’ test taking performance (r(41)=0.57, p=7.1 · 10^-5^) as well as gaze position (r(38)=0.52, p=5.6· 10^-5^), and pupil size (r(39)=0.52, p=6.7· 10^-4^). To a lesser extent this is also true for heart rate (r(41)=0.30, p=0.05). In contrast, ISC of respiration does not correlate with memory performance (r(41)=0.01, p=0.94).

### Correlation with others depends on the individual’s attention to the stimulus

The interpretation that cognitive processing of a common stimuli is required for ISC to emerge is consistent with the observation that it is modulated by attention for many of these modalities. Specifically, when subjects are distracted from the stimulus, ISC drops significantly for gaze position, pupil size, EEG as well as heart rate ^3,10,19^. To compare these effects across modalities and determine their time scales, we performed a new experiment in which N=29 subjects watched videos while normally attending, and then again while distracted by a mental arithmetic task. In this Experiment 2, we used 6 instructional videos with a total duration of 31 minutes. We compute ISC again resolved by frequency but separately for the attentive and distracted conditions (Fig. 5A). We find that ISC is significantly weaker in the distracted condition for all modalities (respiration was not measured in this experiment). We also analyze the power spectrum of these signals to determine if the attentional effects are reflected in the dynamic of these signals (Fig. 5B). We find generally weaker effect sizes for the power spectrum (Fig. 5C). This suggests that attention does not strongly affect physiological dynamics, but rather, fluctuations align in time across subjects when attending to the stimulus and not otherwise. (See Supplement for more detailed discussion of these results for HR and EEG).

**Figure 5.**
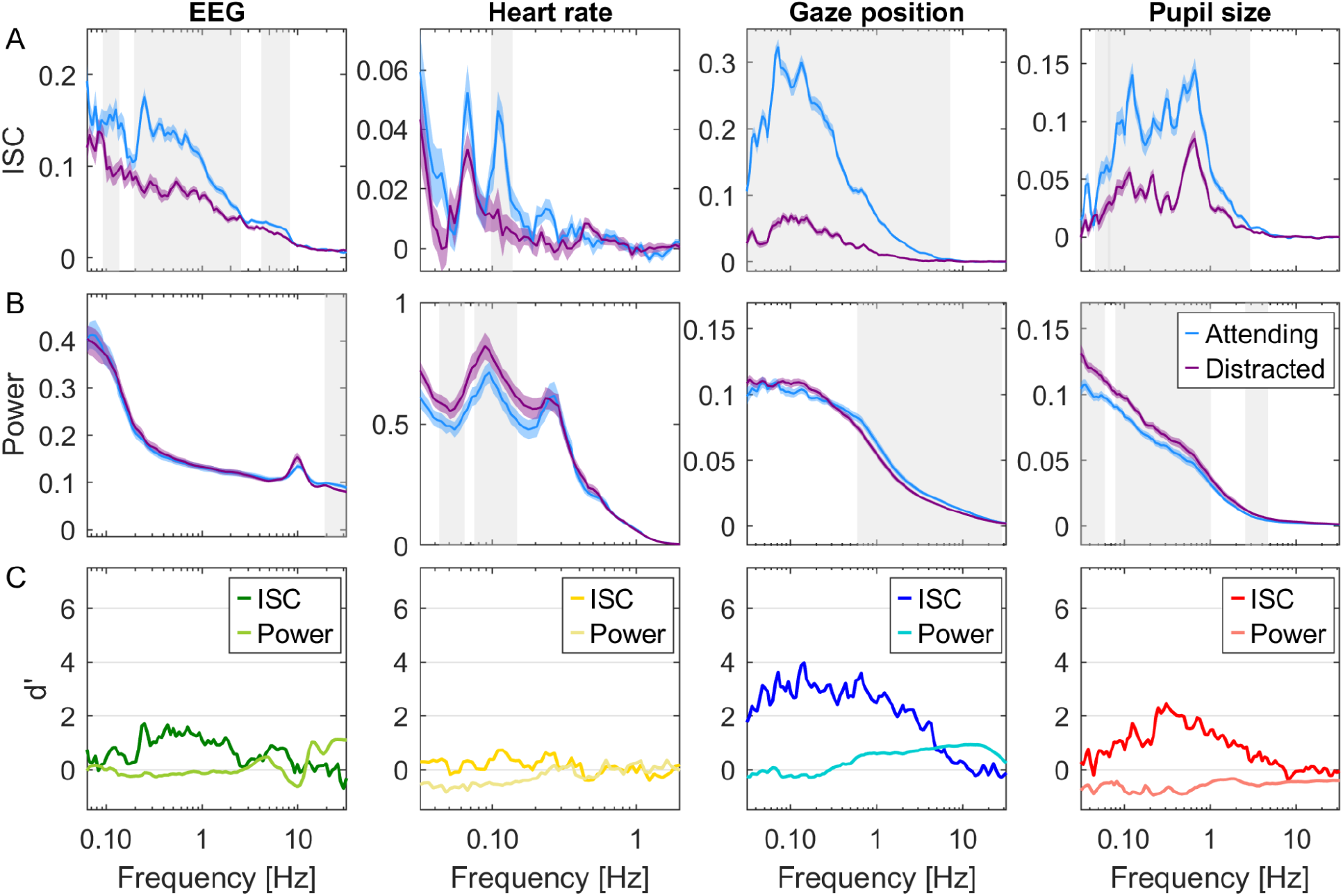
Attentional modulation of ISC and Power resolved by frequency. Here subjects watch instructional videos (31 min total duration, Experiment 2) attentiding normally (blue) or distracted (purple) by performing a mental arithmetic task. **A**: Coherence spectra measure inter-subject correlation resolved by frequency. The color shaded areas indicate standard error of the mean (SEM) across subjects. Significant differences between attending and distracted conditions are established for each band using a t-test, corrected for multiple comparisons with one-dimensional cluster statistics (grey-shaded frequency range indicates p<0.01, N=29, cluster threshold corrected). **B:** Power spectra measure power of signal fluctuations resolved by frequency. Shaded areas as in B. **C:** Effect size is measured as Cohen’s d’ with variance estimated across subjects. ISC and power are computed on band-pass filtered signals (as in Fig. 3.) and averaged over 6 videos.

### ISC occurs only for physiological signals that are coherent with brain signals

Given our hypothesis we predict that significant ISC in the physiological signals occurs if, and only if the signal correlates with brain activity within subjects during video watching. In this view, when brains correlate, so will other physiological signals that are driven by cognitive processing in the brain. The alternative hypothesis is that the brain-body connection itself is modulated by attention for these modalities. To test for this we measured the within-subjects coupling between EEG and heart rate, pupil size and respiration (Fig. 6). Specifically, at each frequency band we extract a component of the EEG that best correlates with the respective signals and report the strength of this correlation, which we call within-subject correlation (WSC). We find that gaze position, pupil size and heart rate significantly correlated with brain signals in some frequency band (Fig. 6, and S3 for distribution over the scalp). This is trivially expected for gaze position due to saccade-evoked potentials ^36^, but this has not been previously reported for pupil or heart rate. Importantly, we find no correlation between brain potentials and respiration. We find only minor differences between the attend and distract conditions in these coherence spectra (for respiration we only have data on the attentive condition). The picture that emerges therefore is that the brain correlates with other subjects when it engages with the stimulus, and that this carries over to physiological signals as a result of an endogenous brain-body coupling (Fig. 8), which is relatively stable with regards to attentional state during passive video watching.

**Figure 6:**
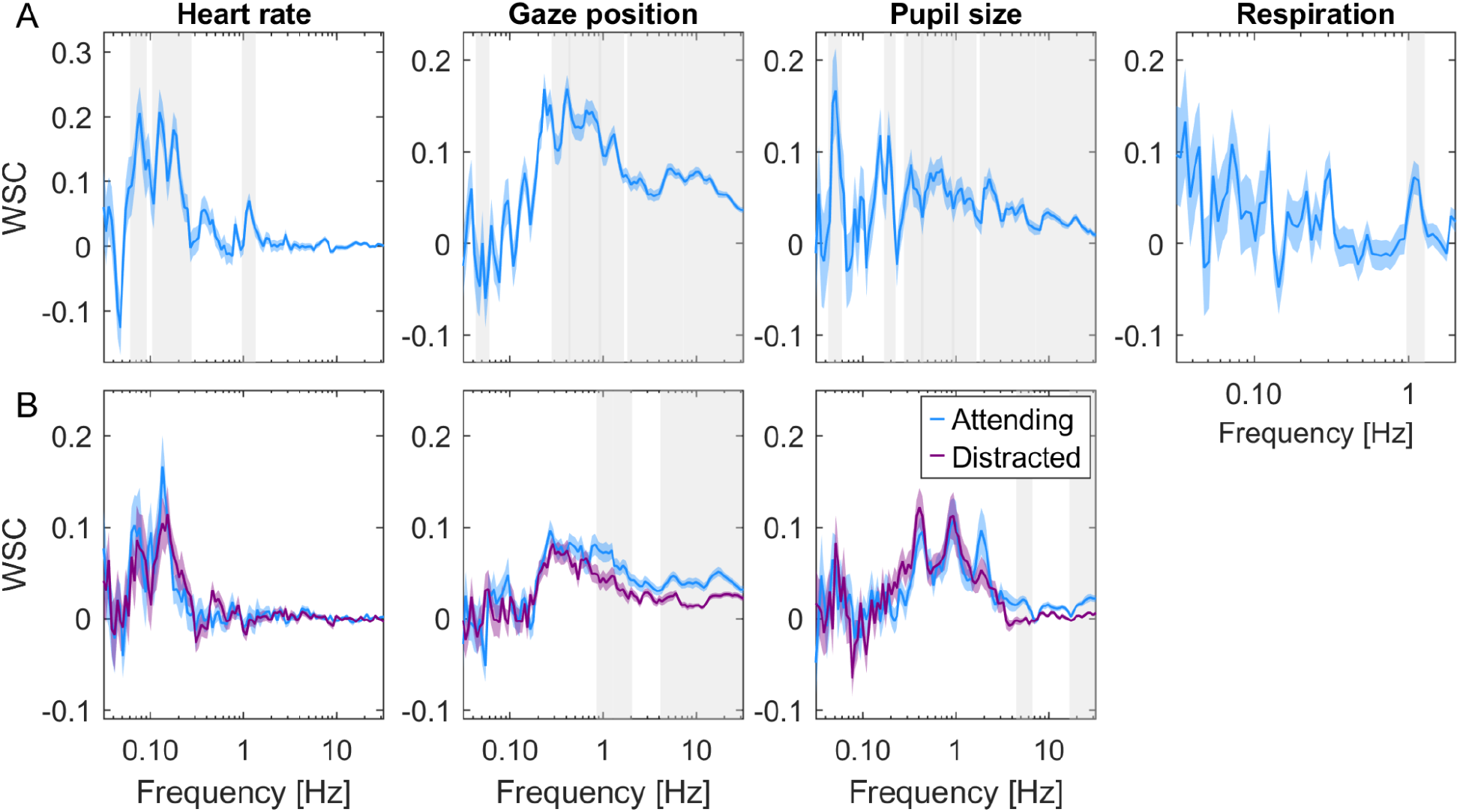
Within-subject coupling between EEG and other signal modalities during viewing of videos. Within-subject correlation between EEG and gaze position, pupil size, heart rate and respiration. The within-subject correlation is computed as the correlation between each signal modality and spatially filtered EEG. The spatial filters are found as the least square fit of scalp electrode potentials with the respective signal modality (computed separately for each frequency band; see supplement for variation across bands). **A:** WSC computed by combining the 3 stimuli used in Experiment 1. Blue-shaded area indicates SEM over subjects. Gray-shaded frequency-band indicates significant difference from zero (p<0.01, N=92 subjects, cluster corrected t-test on test data). **B:** WSC values are computed separately for the attentive and distracted viewing conditions. WSC values are the average across the 5 stimuli used in Experiment 2 with N=29 subjects, and the shaded area around the average WSC values is the SEM over subjects. Gray-shaded frequency range indicates a significant difference between attending and distracted conditions (p<0.01, paired t-test N=29, cluster corrected).

**Figure 7:**
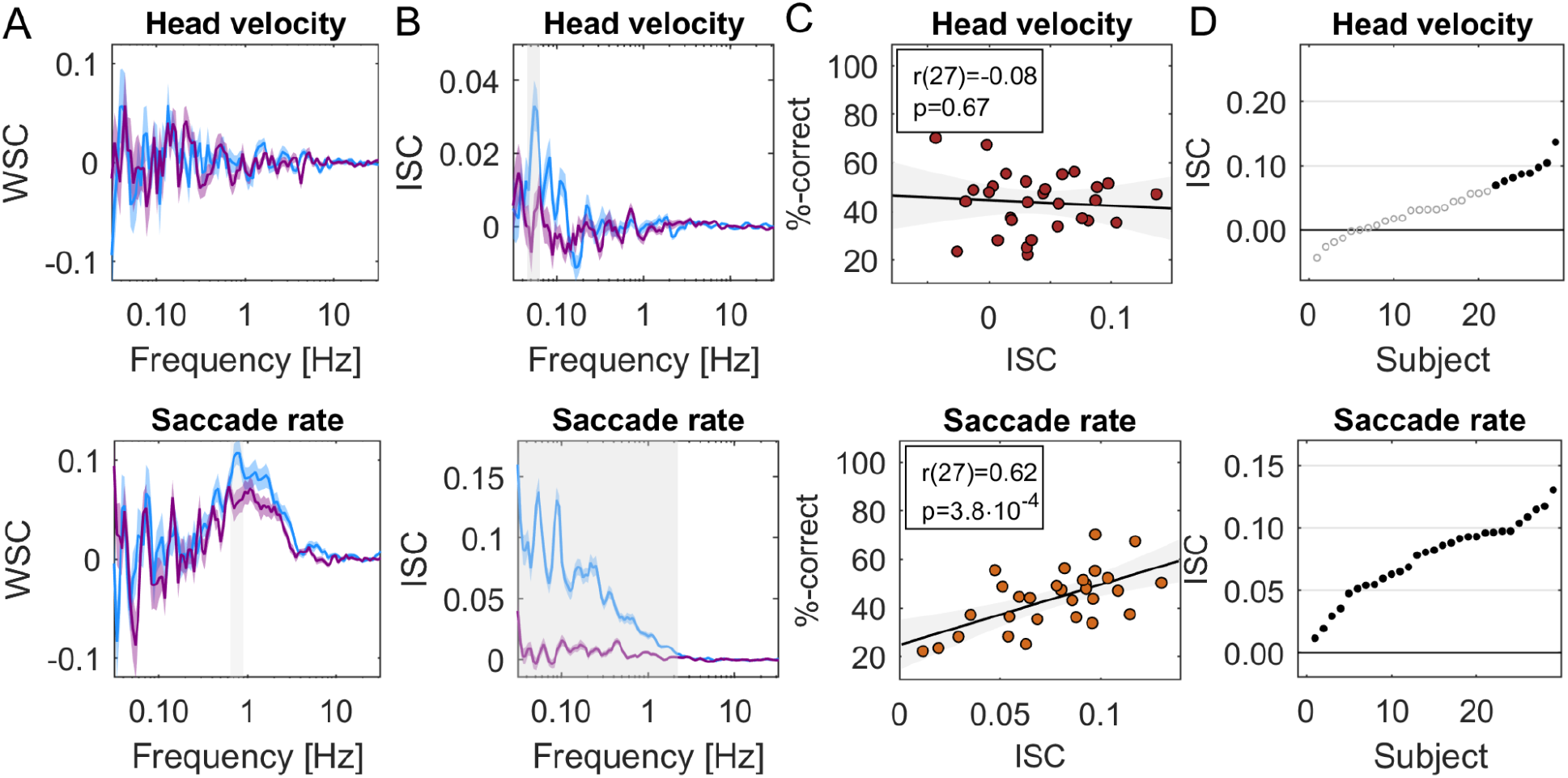
Predictions and test for saccade rate and head velocity. Here using data from Experiment 2 from 29 subjects we test predictions on saccade rate and head velocity. **A:** EEG has significant within subject correlation (WSC) with saccade rate but not head velocity, and neither difference with attentional state (attend: blue, distract: purple). **B:** Significant ISC is observed for all subjects for saccade rate (29/29) but only a subset of subjects for head velocity (8/29). **C:** Saccade rate, but not head velocity are predictive of memory performance across subjects. **D:** Attention modulates ISC of saccade rate but not head velocity. Saccade rate is computed similarly to heart rate. Head velocity is calculated as the magnitude of the Hilbert transform of head position, combining horizontal, vertical and depth directions with Euclidean norm. Significance established using the same methods as above.

**Figure 8:**
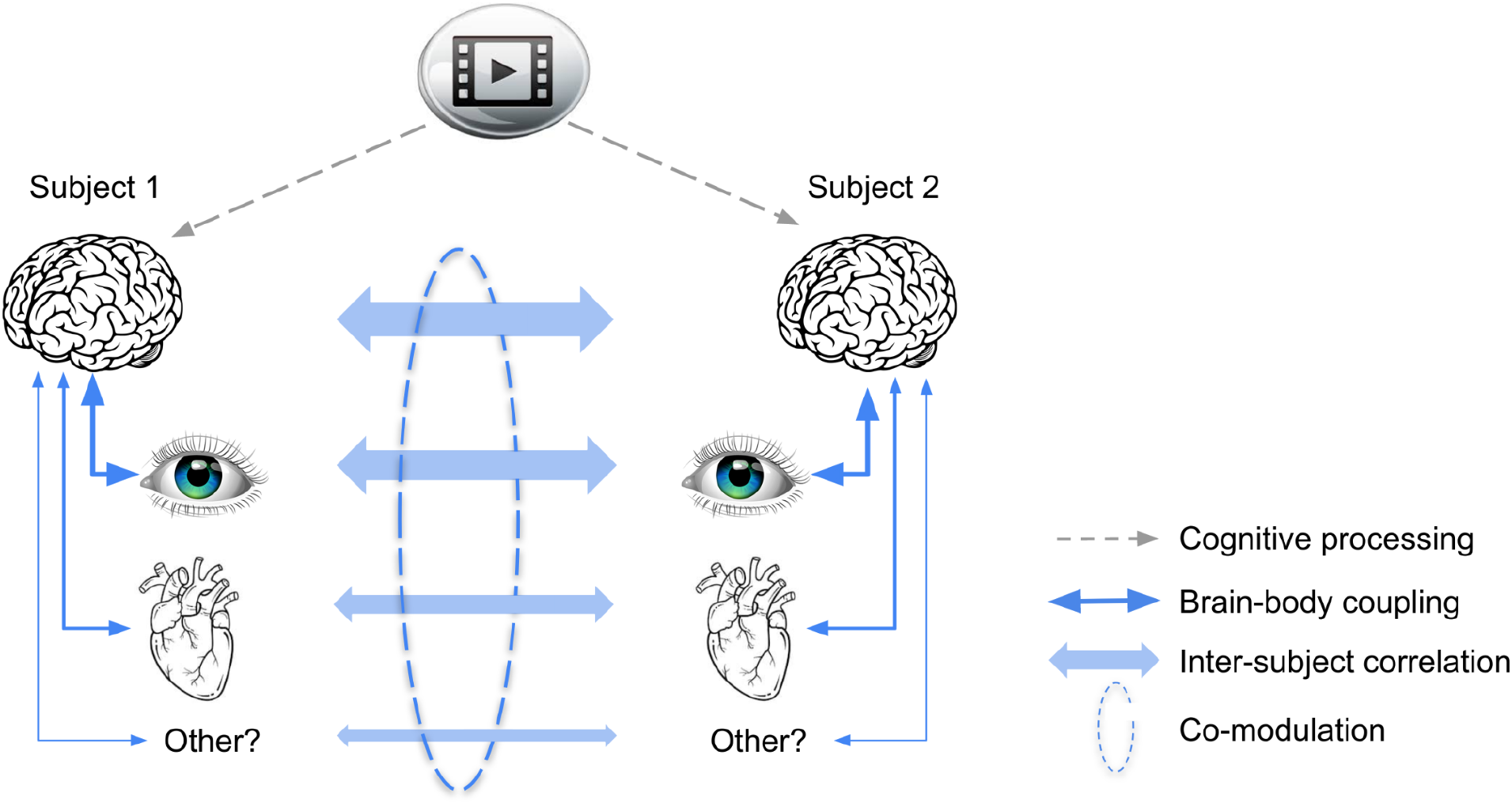
Schematic summary of results and proposed common mechanism. A video stimulus is processed cognitively by the brain, similarly in different subjects (dashed arrows -- indicates cognitive processing encompassing perception and cognition; obviously signals enter the brain through eyes and ears). This cognitive processing causes similar fluctuations in signals of the body that exhibit robust brain-body coupling (solid arrows). Therefore, brain, physiological and behavioral signals correlate between subjects (bold arrows). Subjects’ attention variably engages with the stimulus, such that cognitive processing as a mediator of common fluctuations varies, across subjects and time. This modulation results in a co-modulation of ISC in different signals. Links indicated in blue have been measured in this study as correlation. Narrow arrows are hypothesized causal effects (note that brain-body links may be bidirectional, which is obvious for the eyes and is discussed below for the heart as well).

### Brain-body connection predicts inter-subject correlation for novel signals

Our theory therefore is that signals correlate between subjects, but only if there is a robust coupling between brain signals and the physiological or behavioral signal in question (Fig. 7A). We tested this on two additional signal modalities that have independently been proposed as markers of arousal. One is head velocity^37^, following the basic notion of arousal as movement of the body. The other is saccade rate, which has been recently linked to effort and task engagement ^38^. First we measure if either are coupled to EEG (on data from Experiment 2), and find this to be the case for saccade rate but not head velocity. We therefore predict that saccade rate and not head velocity will be correlated across subjects and this correlation will be modulated by attention and predict memory. These predictions are all confirmed by the subsequent analysis (Fig. 7B and 7C), and indeed, co-modulation of ISC between all modalities across subjects (Fig. S1) and across time (Fig. S2) is consistent with this theory. Note that correlated gaze position does not necessarily imply correlated saccade rate, as it only captures the number of saccade per unit time. On the other hand, head velocity could have been expected to correlate as large saccades tend to be accompanied by corresponding reorienting of the head ^39^. Therefore, these positive and negative examples were not trivially anticipated without the proposed theory.

### Conventional indices of arousal and reproducibility of effects

A more specific hypothesis to explain inter-subject correlation or heart rate and pupil size may be that the video causes fluctuations of arousal which are similar across different subjects. To test this we used traditional measures of arousal such as average heart rate (HR) and heart rate variability (HRV), as well as average pupil size, and variation in pupil size. Conventionally, measures of heart rate and pupil are taken on a time scale of minutes, so we computed them for the duration of the videos and averaged cross videos. Overall the results are mixed (Fig. S4): Average pupil size is correlated with ISC of pupil size for Experiment 3 but this does not reproduce for Experiment 2 (Fig. S4A) and we see a similar pattern with pupil variability. For heart rate and heart rate variability we do not see any significant correlation with ISC in any experiment. Pupil size was modulated by attention (Fig. S4B) but did not predict memory performance in either Experiment 2 or 3 (Fig S4C). Heart rate variability was consistently higher in the distracted condition in both experiments tested (Fig. S4E), which is also consistent with the power spectrum of heart rate (Fig. 5B). Heart rate variability was negatively correlated with memory performance but this was not a robust effect across Experiments 2 and 3 (Fig. S4F). In contrast, the correlation of ISC with memory performance reproduced well for all modalities (Fig. S5). Similarly, the modulation of ISC with attention was replicated for all modalities in Experiment 3 (Fig S6). In view of the robust effects with ISC of pupil size and heart rate, these results indicate that these signals are modulated similarly across subjects on a time scale of 10s or less, whereas for longer time scales of minutes, pupil and heart rate are not meaningfully modulated in the present experimental protocol, except for heart rate variability.

## Discussion

To summarize, during viewing of informative videos we found significant inter-subject correlation in all physiological signals that exhibited robust coupling with the EEG. This was the case for heart rate, pupil size, gaze position, saccade rate, but not respiration or head velocity. This is consistent with the theory that correlation between individuals is the result of cognitive processing of a shared stimulus but will only be observed for those physiological or behavioral signals that exhibit robust coupling with brain activity (Fig. 8). Importantly, the strength of this correlation for different signals was co-modulated across subjects, consistent with this theory. This common modulation was predictive of an individual’s memory performance and was attenuated when subjects were distracted from the stimulus.

The strength of ISC observed here as well as the modulation with attention and correlation with memory performances are in line with previous reports for heart rate^10^, gaze position, pupil size ^3^ and EEG ^17,19,34^. The novel finding is that ISC is co-modulated in these signals across subjects and across time. We provide a novel theory as to which signals do or do not correlate and confirm this with saccade rate and head velocity as positive and negative examples, which had not been previously reported in the literature.

The co-modulation observed here is surprising given the diversity of factors that have been previously proposed to underlie interpersonal physiological synchrony. Leaving out brain signals, this includes emotions (HR ^5–7,11^), arousal (HR ^4,24^, head velocity ^37^), empathy (HR ^6,8,23^, skin conductance ^22^), attention (eye movement ^3^, HR ^24,25^), auditory features (HR and respiration ^41,42^) visual features (gaze position ^43,44^), physical movement (respiration ^45,46^, HR ^21^), social interaction (HR ^47^), and more. We argue more generally, that cognitive processing of the stimulus is a minimum requirement to evoke any of these constructs in a viewer’s mind, and indeed, it suffices to explain the attention and memory effect we observed, namely, without processing the information imparted in the stimulus it is not possible to recall this information, and distracting the viewer will disrupt processing and thus reduce inter-subject correlation. If interpersonal synchronization was really the result of the various factors as postulated in prior literature, it is not clear how this diversity of factors could have affected different modalities similarly, and even less that they should have modulated ISC in unison across time and across subjects. That said, the weak co-variation observed across time suggests that a diversity of factors may drive signal fluctuations differentially in each modality.

An alternative interpretation of the present results is that low-level stimulus features are the source of co-variation in ISC. For example, visual dynamics may attract similar eye movement and this affects ISC of brain signals. With the strength of visual dynamics itself varying over time this could cause a co-modulation of ISC of the eyes and brain. There may be any number of stimulus features that guide visual exploration and could cause such co-variation in a bottom-up manner. However, this alternative interpretation does not explain the co-modulation of ISC observed across subjects. It also does not readily explain the co-modulation in heart and eyes, nor the observation that ISC of the heart is predictive of memory, a phenomenon that persists for audio-only narratives ^40^. In contrast, the effects of attention and on memory observed here are straightforward consequences of the cognitive processing hypothesis, which in our definition includes perceptual processing of the stimulus.

So what might be the common factor that affects cognitive processing of the stimulus, and co-modulates the strength of inter-subject correlation? We speculate that it is related to engagement with the stimulus. In previous work with EEG during passive exposure to narrative audio and video we introduced the concept of attentional engagement with the stimulus ^13,17,34,48^. We demonstrated that an objective behavioral measure of stimulus engagement correlates with ISC of the EEG measured on times scales of 10 seconds ^49^, presumably reflecting fluctuations of attention^19^. In the vision literature “attentional engagement” implies not only attracting attention (gaze) but also processing features of the target of attention ^50–53^. The notion that the ISC of EEG reflects a level of engagement with the stimulus has been adopted by a number of groups ^20,54,55^. The new finding here is that this engagement may be a common factor that also modulates the strength of fluctuations in heart rate, pupil size, saccade rate and gaze position, all in unison. We find this co-modulation here across subjects and across time.

Spontaneous fluctuations in fMRI signals have been linked to EEG measures of arousal or alertness on similar time scales ^56,57^. There is also evidence that fluctuations in functional connectivity of fMRI correlate with HRV^58^ and can be used to measure attentional state on a time scale of minutes^59^. Activity of the “default-mode network” is more correlated across subjects when they report to be more engaged with the stimulus ^60^ and it is increased in more memorable moments of the narrative^16^. The precuneus, which is part of the default-mode network, shows elevated activity during periods of high ISC of the EEG^13^. Whether modulation of inter-subject correlation in other physiological signals is indeed related to the task engagement states observed in fMRI will have to be determined in future work by simultaneously recording fMRI along with other modalities.

Regardless of the interpretation, this co-modulation of physiological responses is surprising for another reason -- the dynamics of each of these signal fluctuations and their underlying neural control are quite diverse. For instance, neural control of HR and pupil size are ascribed to entirely different brain structures. While pupil dilation has been most closely linked to activity in superior colliculus and locus coeruleus ^28^, heart rate is controlled by midbrain structures that are modulated by input from the amygdala, cingulate and insula ^26^, although the LC also projects to nuclei involved in cardiac regulation ^61^. Additionally, the dynamics of these signals, as we have seen in their power spectra, differ considerably between modalities and the attentional effect on ISC manifests in different frequency bands for each. We note that the coupling between EEG and each of the modalities did not differ significantly with attentional state. Note also that the effects of attention on the power spectrum were generally small. Therefore, cognitive processing does most likely not substantially change the ongoing endogenous physiological dynamic within subjects. Instead, it appears to only time-align the existing dynamic to the external stimulus.

Our theory led us to prospectively analyze the correlation between EEG and various physiological signals. While a correlation of EEG with gaze position is well established (in fact, it is considered a nuisance artefact ^62^), to our knowledge this is the first study to report a correlation of EEG with heart rate, pupil size and saccate rate. The theory also correctly predicted that saccade rate would correlate across subjects, something that has not been reported previously in the literature. Even less obvious was that this correlation of saccade rate would be modulated by attention and predict subsequent memory performance.

We did not find ISC in breathing, which one might have expected. For instance, in moments of suspense, viewers might be “holding their breath” and thus changing the dynamic of heart rate, which is closely linked to breathing ^63^. Breathing seems to align to the timing of a behavioral task ^64^, but here participants were not asked to do anything specific during viewing. Generally, studies that show correlated breathing involve some concurrent physical activity such as singing ^46^, speaking ^65^ or dancing ^45^ and correlation is not observed during passive listening to speech ^66^, though some effects have been found when listening to music^41^.

The fact that gaze positions are robustly correlated between subjects might not come as a surprise since eye movements are heavily driven by the dynamic of visual stimuli ^67^. We deliberately chose stimuli with compelling visual dynamics, such as animated graphics. The pupil size is known to be affected by luminance, but also higher level factors such as problem solving and cognitive effort ^30,68^, affective processing ^32,69^ and attention ^70^. So to see the pupil size correlated across subjects might be expected. That this pupil correlation should be modulated by attentional engagement is consistent with the argument that task engagement is reflected in pupil dynamics on similarly short time scales^38^. That the ISC of pupil should be co-modulated with ISC of HR was not expected from the literature. We have not analyzed eye blinks, however, given reports of a correlation of blinking across subjects during movies ^71^, we very much expect that this will behave similarly to the other measures we have taken from the eyes. Given the coupling that has been observed between EEG and gastric rhythms ^72^ we also predict that these will show significant ISC.

Pupil and heart rate have often been linked to arousal (see below), although the term “arousal” has quite diverging definitions in the literature. We expected that head velocity could serve as a marker of physical arousal ^37^, and thus may also correlate. Correlation may also be expected given the inter-subject correlation of gaze position and the fact that large eye movements tend to be accompanied by corresponding head movements ^39^. We did not find significant ISC in head velocity in our smaller datasets. Consistent with that, ISC of head velocity did not significantly correlate with memory or attention, nor did head velocity significantly correlate with brain potentials. We did, however, find a weak ISC in the larger dataset of 92 subjects (see Supplement), and therefore one might be able to resolve these effects on larger datasets. Indeed, our theory does not stipulate an all-or-nothing effect. A weak brain-body link may lead to similarly weak physiological ISC and related effects.

Correlation of heart rate between subjects has previously been linked to emotional processing of video stimuli ^8,9^, which may not have been a main driving factor in our mostly informative videos. In social contexts, heart rate synchronization has also been linked to empathy ^73^ and social bond ^74^. We propose that cognitive processing of the natural stimulus is a requirement to infer the emotional valence of the film ^9^, for an audience to bond with a performer ^4^ or mother and child to interpret social cues ^47^. Thus cognitive processing of the external natural stimulus is the common denominator, even for interpersonal heart rate synchrony. One caveat to this conclusion is that we did not look for other factors that could have driven the different modalities differently. All videos were designed to communicate factual information related to science and technology, and not, for instance, evoke suspense or emotions ^5,6,11,60^. So it is possible that we did not find other factors simply because we did not manipulate other factors with our experimental paradigm. Future work should consider varying stimulus properties along different dimensions.

There is an abundance of studies linking cognitive factors to heart rate variability and pupil responses. These are typically measured on times scales from many seconds to minutes. In our analysis this is captured in the power spectra, which measure the magnitude of fluctuations of HR and pupil size averaged across minutes. In contrast, ISC captures reliable timing of these fluctuations on short time scales (from 30s to less than a second; i.e. the phase of these fluctuations). We are agnostic as to the specific aspect of cognitive processing that reliably drives these fast fluctuations. Indeed, HRV and pupil dilation have been linked to a great number of different cognitive factors. For instance, HRV has been linked to emotions ^75^, stress ^76^, attention ^77^, empathy ^6^ and other cognitive factors ^27^. Pupil dilation has been linked to arousal ^78,79^, emotions ^32,69^, effort ^79,80^, surprise ^31,81^, task engagement^38^, saliency^82^, attention^83^ and more. As such, these measures are quite sensitive, but also quite unspecific. We did find an increase in power (i.e. HRV) when subjects are distracted from the videos by a secondary task. Previous studies show that reduced HRV is linked to hyper-vigilance and less flexible attentional control ^77^. For the pupil, power captures the magnitude of pupillary response. Previous studies find a pupillary response with shifts in attention ^70^. Here, we found an increase in power when viewers are distracted from the stimulus by a secondary mental task. In our view, the traditional measures of heart rate and pupil size variability may be quite sensitive to the specifics of the experimental manipulation. In contrast, the ISC measure gains in robustness by being agnostic to the specific factors elicited by the stimulus. Any of the many factors that might cause a rapid fluctuation in physiological signals will contribute to ISC.

The cognitive control of heart rate is a crucial prerequisite for the co-modulation observed here. Indeed, central control is well established and explains the effects of cognition on heart rate and HR variability ^26,27^. For instance, heart rate variability correlates with neural activity measured with fMRI ^76^. There is also evidence for the opposite causal direction, whereby volitional control of breathing affects heart rate and in turn neural activity ^84^. The heart beat itself seems to affect saccade timing^85^, evoke EEG ^86^, as well as visual detection performance^87^. Also, strong stressors can increase heart rate and at the same time enhance the magnitude of evoked responses in EEG ^88^. But none of these studies report a link between EEG and heart as we have found here, nor do they anticipate that ISC is co-modulated in these modalities.

A recent theory proposes a common drive to pupils and saccades ^38^. The theory posits that this common drive is affected by a variety of cognitive processes, in particular “task engagement”, but authors distinguish this from “arousal”. This common drive is postulated to be the activity in superior colliculus (SC). Indeed, the SC along with the locus coeruleus (LC) are midbrain nuclei that mediate cognitive effects on the pupil ^28,82^. A common drive of pupil and saccades reconciles the dependence of pupil dilations on arousal and effort as well as its fluctuations during a task. Consistent with this we find that pupil size and saccade rate both behave similarly with attention and memory and importantly the ISC in these two modalities are co-modulated. Note that the LC also connects to midbrain structures that control cardiac function^89^ and may mediate cognitive effects on heart rate^61^. Therefore, the LC is well situated to mediate cognitive effects on pupil size, saccades rate and heart rate.

In conclusion, this work brings together two separate research fields which have demonstrated correlation between subjects in ecologically valid scenarios, namely, interpersonal physiological synchrony ^1^ and inter-subject correlation of neural activity ^12^. We postulate that cognitive processing of a natural dynamic stimulus is required to drive and coordinate behavioral responses of the body. Our rationale is that humans have evolved to quickly respond to the natural stimuli in their environment, and to do so, the brain processes information and prepares the body to react. This leads to rapid, but reliable fluctuations of heart rate, pupil size, gaze position, etc. on the scale of seconds. If the brain is not processing the stimulus effectively, i.e. similarly across subjects, then the body will not respond appropriately and these fluctuations will no longer be guided by the stimulus. As the brain changes in attentional state on time scales of 10s or longer the correlation induced across subjects is modulated for different modalities in unison. Although our study here is only correlational, we do postulate a causal effect of cognitive processing on physiological and behavioral responses that is similar across subjects and modalities. Evidently there could be a bidirectional interaction. This is most obvious for eye movements, as our view can affect our cognitive state. A bottom-up effect has also been hypothesized for heart signals ^84^ and is the basis for some meditation practices that focus on breathing ^90^. Establishing a causal direction of these effects will be difficult but is a worthwhile topic for future research.

## Materials and Methods

### Participants

158 subjects participated in one of 3 experiments, where they watched informative videos. In Experiment 1, N=96 participated (Female 51, age 18-49, M=25.33, SD=7.29; 4 subjects were removed due to bad timing or bad signal quality). In Experiment 2, N=32 subjects participated (21 females, age 18-57, M=25.93, SD 8.94 years; 3 subjects were removed due to bad signal quality). Lastly in Experiment 3, N=31 participated (19 females, age 18-50, M=25.79, SD=8.13 years; 2 subjects were removed due to bad signal quality).

### Stimuli

A complete list of the video stimuli is provided in Table S1 of the Supplement. These stimuli have been used in previous studies ^10,34,91^. For Experiments 1 we selected three videos from YouTube channels that post short informal instructional videos namely ‘Kurzgesagt – In a Nutshell’ and ‘Minute Physics’. The videos covered topics related to physics and biology with a short duration (Range: 3-6.5 minutes, total duration 15.5 minutes). In Experiment 2 we selected 6 videos from ‘Khan Academy’, ‘eHow’, ‘Its ok to be smart’ and ‘SciShow’ which are popular online educational channels on YouTube. The videos covered topics related to biology and physics, with a short duration (Range: 4.5-6.5 minutes, total duration 31 minutes). In Experiments 3 we selected 5 informal instructional videos again from YouTube, covering topics related to physics, biology, and computer science with a short duration (Range: 2.4 – 6.5 minutes, Average: 4.1 +- 2.0 minutes). Data on gaze position and pupil size for Experiment 2 have been previously analyzed^3^, as well as data on heart rate from Experiment 3^10^. All other data and analyses are new to this study, i.e. all of the data from Experiment 1, EEG, heart rate, saccade rate and head velocity from Experiment 2, and EEG, pupil size, gaze position, saccade rate and head velocity from Experiment 3.

### Procedure

All experiments were carried out at the City College of New York with approval from the Institutional Review Boards of the City University of New York. Documented informed consent was obtained from all subjects at the start of the experiment. Subject wore headphones and watched the videos on a 19” monitor while seated comfortably in a sound-attenuated booth. In Experiment 1, subjects watched 3 instructional videos while their electro-encephalogram (EEG), electro-oculogram (EOG), electro-cardiogram (ECG), respiration, pupillary responses, gaze and head position were recorded. In the second experiment subjects watched 6 instructional videos while their EEG, ECG, pupillary responses, gaze and head position were recorded.

In all the experiments, subjects were instructed to watch the videos normally as they would at home, while being relaxed and sitting still. We refer to this as the attentive conditions (A). After they had watched the videos, subjects were given a four-alternative forced-choice questionnaire covering factual information imparted during the video (11 – 12 recognition questions per video; see Table S1 of the Supplement). The videos and question pairs were presented in random order. In Experiment 1, subjects were not aware that they would be tested on the material, whereas in Experiments 2 and 3 the test was anticipated. After answering questions, in Experiments 2 and 3 subjects were asked to watch the videos again, but this time to silently count in their mind backwards in steps of 7 (starting from a prime number picked at random between 800 and 1000). The second viewing with concurrent counting is referred to as the distracted condition (D).

For segmentation of the physiological signals we used common onset and offset triggers, in addition a flash and beep sound was embedded right before and after each video which were recorded using a StimTracker (Cedrus) to ensure precise alignment across all subjects. To enable all modalities to be on the same timescale we used the lab streaming layer protocol. In addition we used a linear regression model to convert timestamp between each modality estimated using the common triggers.

### Recording and preprocessing of EEG

The EEG was recorded at a sampling frequency of 2048 Hz using a BioSemi Active Two system. Participants were fitted with a standard, 64-electrode cap following the international 10/10 system. In addition the electrooculogram (EOG) was recorded with six auxiliary electrodes (one located dorsally, ventrally, and laterally to each eye). The EEG and EOG data is band-pass filtered between 0.016 and 250 Hz by the Active two system prior to sampling. The signal was then digitally high-pass filtered (0.5 Hz cutoff) and notch filtered at 60 Hz to remove line noise. To remove artifacts and outliers Robust PCA was used, and subsequently the signal was low-pass filtered (64 Hz cutoff) and down-sampled to 128 Hz. Bad electrode channels were identified manually and replaced with interpolated channels. The interpolation was performed using the 3D cartesian coordinates from the electrode cap projected onto a plane using all surrounding “good” electrodes. The EOG channels were used to remove eye-movement artifacts by linearly regressing them from the EEG channels, i.e. least-squares noise cancellation. In each EEG channel, additional outlier samples were identified as values exceeding 4 times the distance between the 25th and the 75th quartile of the median-centered signal, and samples 40 ms before and after such outliers were replaced with interpolated samples using neighboring electrodes.

### Recording and preprocessing of ECG

The ECG signal was recorded using two ECG electrodes placed below the left collar bone and one on the left lumbar region with a BioSemi Active Two system at a sampling frequency of 2048Hz. The ECG signal was detrended using a high-pass filter (0.5 Hz cutoff) and subsequently notch filtered at 60 Hz to remove line noise. Peaks in the ECG corresponding to the R-waves were found using *findpeaks* (built-in matlab function). The instantaneous HR is computed as the inverse of time intervals between each R-wave. To ensure the same sampling frequency for all subjects this instantaneous HR signal is resampled at a constant sampling rate of 128Hz.

### Recording and preprocessing of gaze position, head velocity and pupil size

Gaze position, head movements and pupil size were recorded using the Eyelink 1000 eye tracker (SR Research Ltd. Ottawa, Canada) with a 35mm lense at a sampling frequency of 500 Hz. Subjects were instructed to sit still while the experiment was carried out, but were free to move their heads, to ensure comfort (no chin rest). A standard 9-point calibration scheme was used utilizing manual verification. Stable pupillary responses were ensured by adjusting the background color of the calibration screen and all instructions presented to the subjects to be the average luminance of all the videos presented during the experiment. After each stimulus presentation a drift-check was performed and the eye tracker was recalibrated if the visual angular error was greater than 2 degrees. Blinks were detected using the algorithm of the eye tracker, and remaining transient artifacts were found using a peak picking algorithm. These artifacts, blinks and 100ms before and after were filled with linearly interpolated values. Head velocity is computed as the absolute value of the analytic signal of the Hilbert transform (root mean square over the 3 direction). Saccades were detected by the algorithm of the eye tracker. Instantaneous saccade rate calculated as the inverse time interval between saccades and upsampled to a constant sampling rate of 2000Hz to match the other eye tracking signals.

### Recording and preprocessing of respiration

The respiration signal was recorded using a SleepSense 1387 Respiration belt placed around the chest of the subject connected to the BioSemi Active Two system recorded at a sampling frequency of 2048Hz. The polarity of the signal was detected using peaks in the respiration signal and inverted to ensure the correct phase of the signal.

### Intersubject correlation analysis of gaze position, pupil size, respiration, heart rate, saccade rate and head velocity

For each of the gaze position, pupil size, respiration, heart rate, saccade rate and head velocity signals, the inter-subject correlation was computed in the following 3 steps: 1. computing the Pearson’s correlation coefficient between a single subject’s signal (each of the six modalities independently) and that of all other subjects while they watched a video. 2. a single ISC value for a subject was obtained by averaging the correlation values between that subject and all other subjects. 3. The two first steps are then repeated for all subjects, resulting in a single ISC value for each subject. For ISC of gaze position we compute the ISC in the horizontal and vertical gaze direction using the procedure as described above separately. To obtain one single ISC_gaze_ value we average the ISC for the horizontal and vertical directions.

### Intersubject correlation of EEG

For the neural signals recorded using EEG the inter-subject correlation was computed using correlated component analysis ^35^. This method finds a model that linearly combines electrodes that capture stimulus-evoked potentials most correlated between subjects. The model consists of several projection vectors that linearly combine electrodes, on which the data is projected (components). The ISC of each component is obtained by computing the correlation coefficients of the projected EEG between each participant and all other participants. We only use components that are significantly correlated above chance (circular shuffle on test set data, see below). This yielded 3-9 components depending which of the three Experiments was analyzed. ISC values are then summed over all significant components.

### Statistical significance of ISC values per subject

To determine whether ISC values are significantly larger than zero (Fig. 1C, 7D), we determine the null distribution of ISC values on surrogate data obtained with circular shuffle statistics^35^. P-value (type I error rate) is then the fraction of shuffles with ISC values larger than in the original unshuffled data. We performed 10,000 circular shuffles estimating p-values down to 0.0001. For EEG, the ISC values are measured on components that have been optimized to provide high ISC values. To avoid an upwards bias in ISC during statistical testing, components are optimized on training data and significance is established for separate test data (two video clips are used for training and one for testing).

### Frequency analysis of intersubject correlation (coherence spectrum)

We performed a frequency analysis to investigate at which time scale the recorded signals correlated between subjects. Each signal from each subject was band-pass filtered using 5th order Butterworth filters with logarithmic spaced center frequencies with a bandwidth of 0.2 of the center frequency. The ISC was computed for each subject in each frequency band for all videos. To obtain a single ISC value per frequency band we average ISC values for all videos and subjects.

### Computation of d-prime (d’) effect of attention on coherence and power spectrum

To compute to which extent both the frequency resolved ISC and Power spectrum is modulated by attention we compute the effect size using d-prime statistics. For each frequency band (see section above) we compute the ISC and power for each subject in the attending and distracted conditions. We quantify the effect size between the two conditions as the difference of the means divided by the standard deviation of the differences (paired Cohen’s d).

### Cluster statistics for difference between attentive and distracted conditions

To determine significant difference in spectra between attending and distracted conditions we use cluster shuffle statistics as follows (for Figs. 5A, 5B, 6B, 7A, 7B). Since different frequency bands in the frequency-resolved ISC (and WSC) are not independent, we use one-dimensional cluster statistics including random shuffles to correct for multiple comparisons following an established procedure ^92^. Briefly, this procedure involves 4 steps: 1) take the difference between the spectrum in the attending and the distracted condition for each subject. 2) compute a one-sample t-test on this difference for each frequency band. 3) clusters are identified as consecutive frequency bands with p-values below 0.01. The t-values within each cluster are then summed. 4) run 10,000 permutations in which we randomly change on half of the subject’s the sign of difference between the spectra computed in step 1. Steps 2-3 are then repeated while keeping the sum of t-values of the largest cluster. Finally, we compare the clusters’ t-values obtained in step 3 with the distribution of permuted cluster t-values obtained in step 4. Clusters with larger than 99 % (corresponding to p-value<0.01) of the pemuted distribution were considered significant after multiple comparison cluster correction. Note that for the contrast in attention all data was used for optimization including attentive and distracted conditions, so that any difference can not be due to the optimization procedure.

### Cluster statistics for significant of inter-subject coherence spectra

To determine if frequency-resolved ISC (and WSC) values are significantly different from zero (Fig. 3A, 3B, 6A) we use a similar cluster statistic as above. For cases involving EEG (correlated component analysis and regression) we use test data to avoid upwards bias due to optimization, which is performed on separate training data. The shuffling and cluster correction procedure consists of the same 4 steps as above, except that we divide subjects in two equal size groups at random. The premise of this is that values around zero will not differ significantly if placed at random in two different groups.^92^

### Within-subject correlation analysis (brain-body coupling)

To determine the coupling between the brain and different signal modalities we compute within-subject correlation between each signal modality and spatially filtered EEG. The spatial filters are found as the least square fit of scalp electrode potentials with the respective signal modality. This is done combining all stimuli but in the case where we differentiate between the attend and distract conditions, we compute spatial filters in each condition separately. For the frequency-resolved WSC, we band-pass filter the signal modality in the same fashion as the time-resolved ISC, namely using 5th order Butterworth filters with logarithmic spaced center frequencies with a bandwidth of 0.2 of the center frequency. To compute the WSC in the frequency resolved condition, we find spatial filters for each band separately. WSC in all instances was computed on test data using leave-one-subject out cross-validation (i.e. regression parameters are estimated including all subjects except for the test subject; WSC is then computed on that test subject; this train-test process is then repeated for all subjects.) Statistical significance is established in the same way as for ISC as described in the two previous sections.

### Common factor analysis of ISC

To find the common factor of ISC between signal modalities we first remove outliers of the ISC data that are larger or smaller than 4 times the interquartile difference of the data. This was done since the standardization of the data is sensitive to extreme outliers. We then standardize the ISC (zero mean and unit variance) and compute the principal components.

## Acknowledgement

This work was funded by the National Science Foundation through grant numbers DRL-1660548.

## Author Contribution

J.M and L.C.P designed the experiments. J.M collected the data. J.M and L.C.P analyzed the data. L.C.P and J.M wrote the manuscript.

## Data availability

Data and code to produce all figures in the paper will be made publically available on Open Source Framework (OSF) upon final publication.

## Supplement

### Stimuli

**Table S1.**
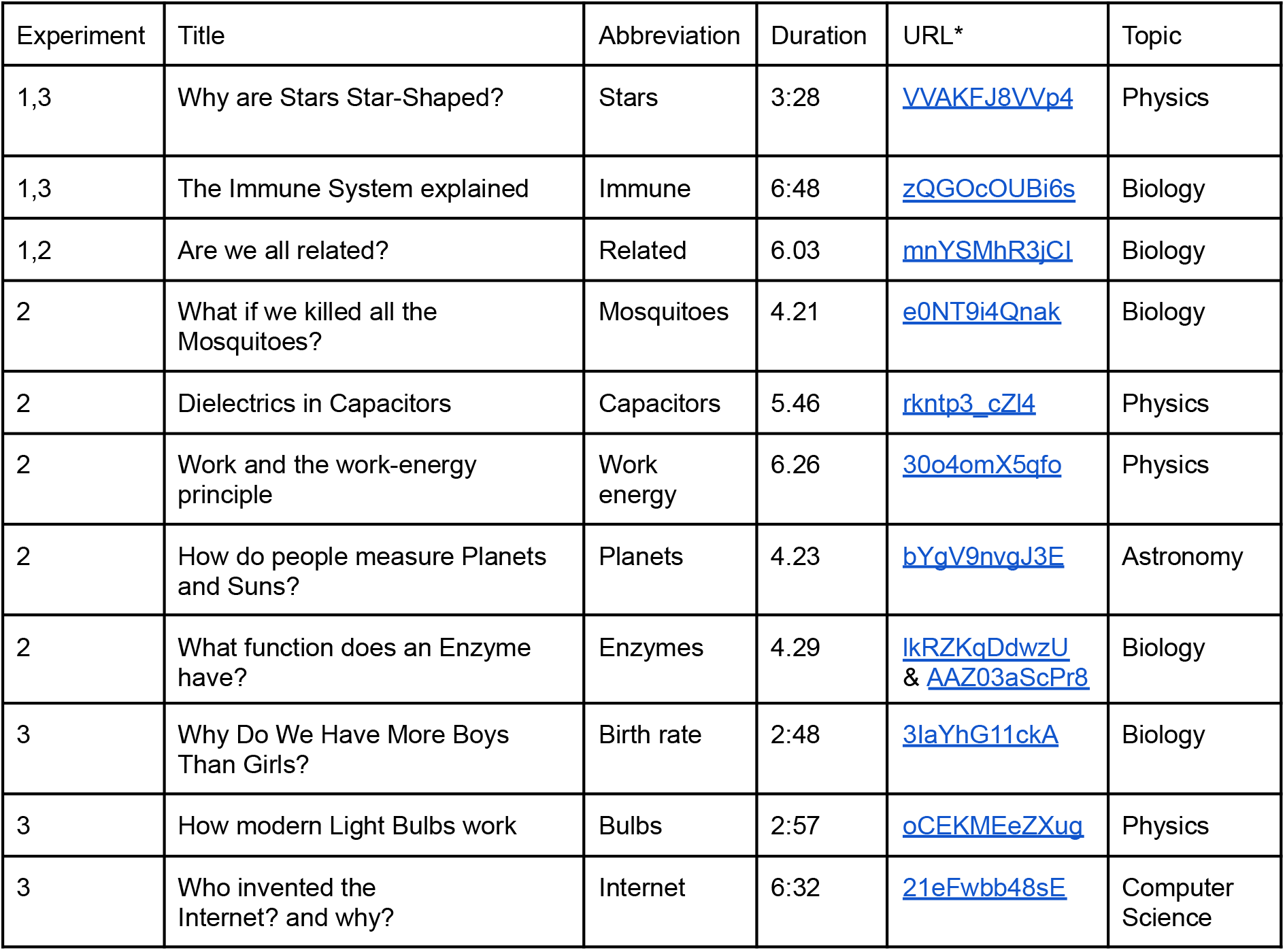
Experiment, title, abbreviation, duration, web address. *URL of videos beginning with https://www.youtube.com/watch?v=

Full list of questions and answer options can be found [here](https://docs.google.com/spreadsheets/d/1cFMGBNyS2MHBTZeUkE6mH8SooZ5Sd-jACK19W3TwRbU/edit?usp=sharing).

**Figure S1:**
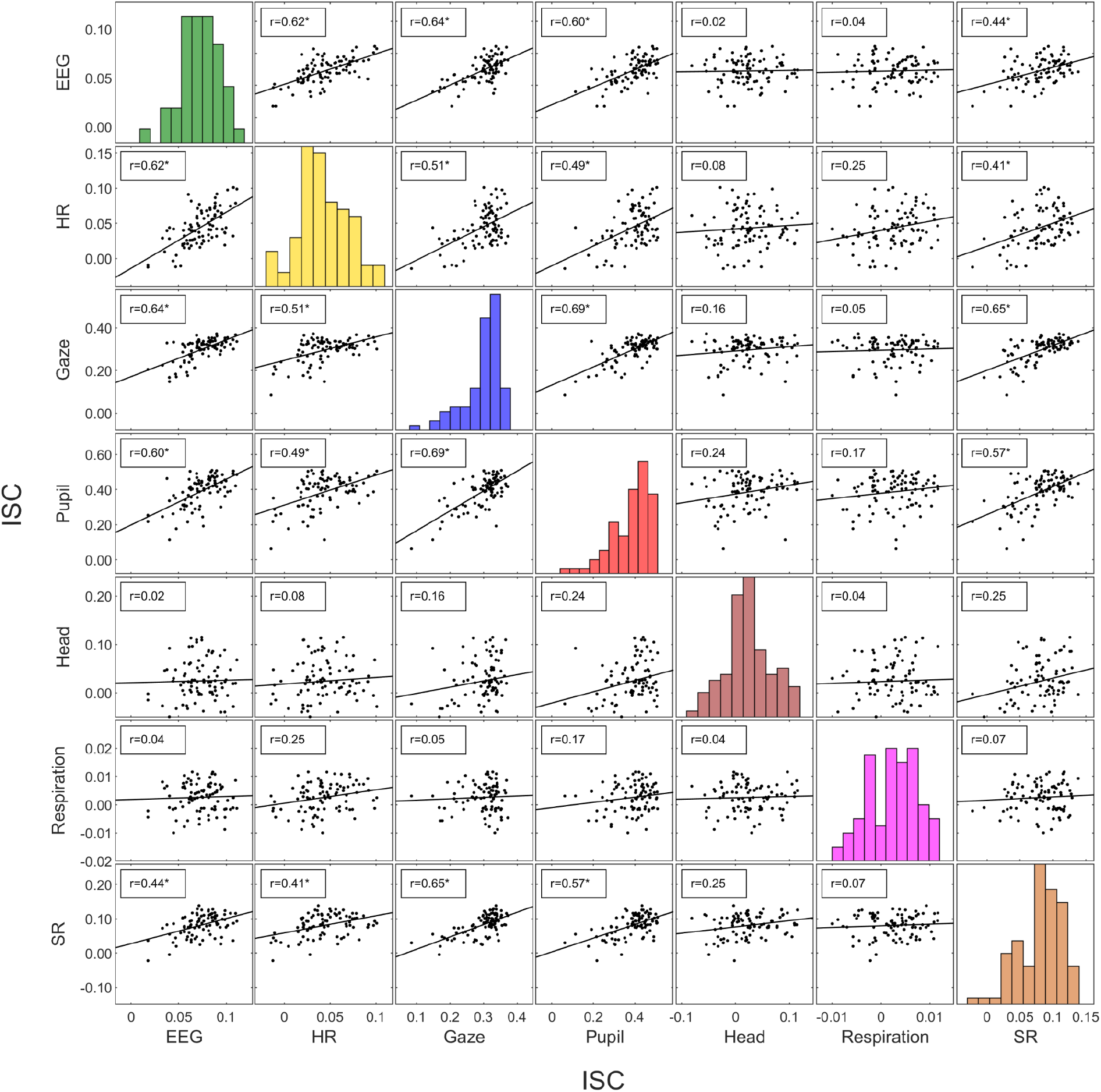
ISC of different modalities is co-modulated across subjects. ISC is computed here for each subject as in Fig 1 (Experiment 1). Here we are including head velocity (Head) and saccade rate (SR), modalities which we tested only after making a prediction based on Fig. 7.

**Figure S2:**
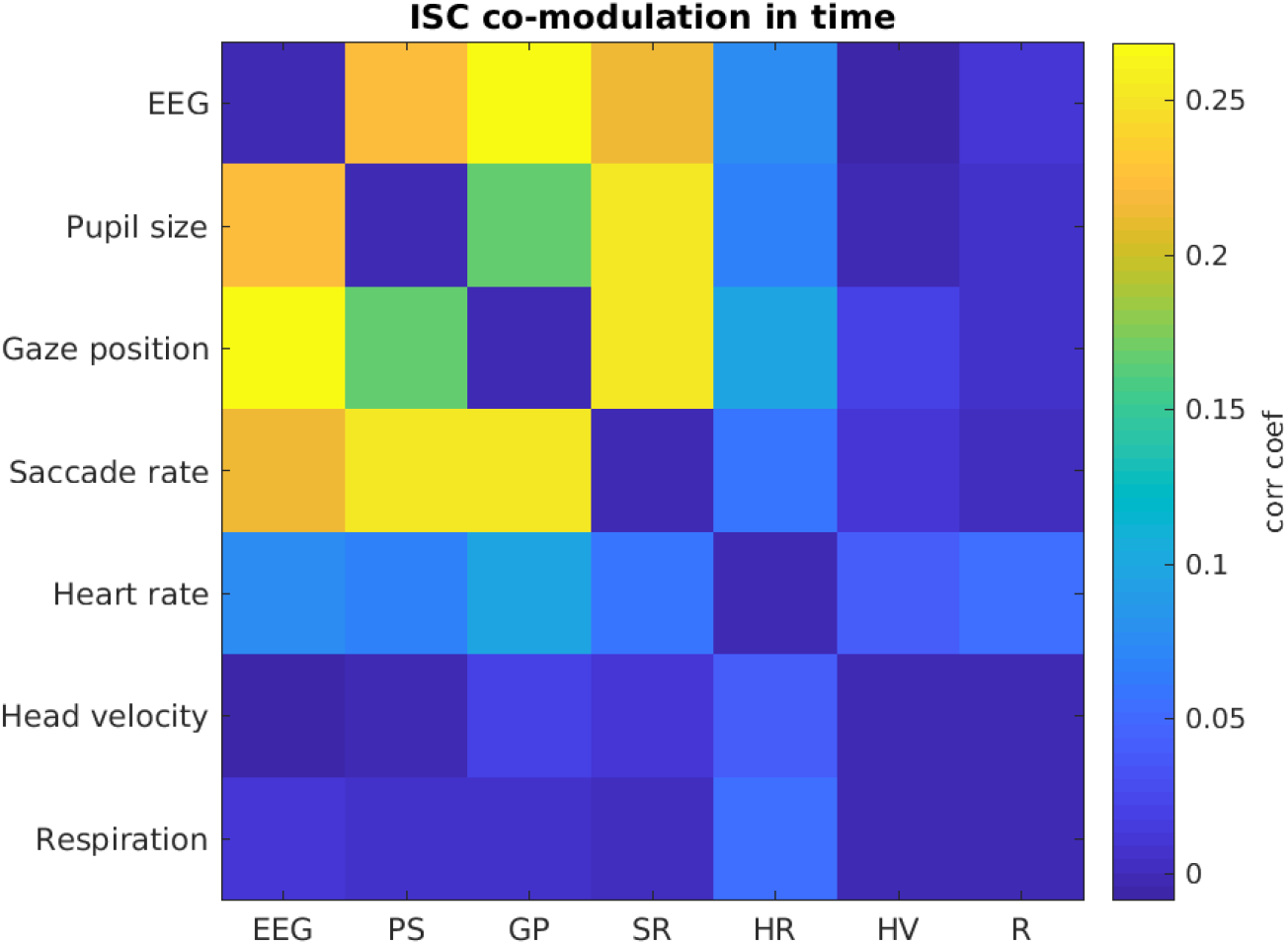
ISC of different modalities correlates weakly across time. ISC is computed here within a sliding time window of 10s width, and correlated between modalities across these time windows (3 video clips lasting 9 min 14 sec) and averaged across subjects (N=43) from Experiment 1. Statistical analysis for a few individual cells is in Fig 3C. Here we are including head velocity and saccade rate, modalities which were tested only after making a prediction based on Fig. 7.

### Discussion of Figure 5 for EEG and HR

For EEG, the coherence spectrum (frequency-resolved ISC, Fig. 5A, first column) is dominant in low frequencies (down to the lowest frequency we measured) and decays substantially once we reach 10Hz. The predominant 10Hz peak in the power spectrum (Fig. 5B, first column) is known as “alpha” activity, and does not appear to correlate across subjects.

For HR, the coherence spectrum (Fig. 5A, second column) has two pronounced peaks that fall within the 0.1 Hz band in the HR power spectrum (Fig. 5B, second column). This band (0.09-0.14Hz) characterized as “low-frequency” in the literature on HR variability ^93^ is strongly modulated by attention in the present study. Interestingly, this peak is known to be attenuated also during slow-wave sleep.^94^ The “high frequency” band (around 0.3Hz) in the power spectrum does not appear to correlate across subjects. This band corresponds to the dominant respiration frequency (see Fig. 3).

**Figure S3:**
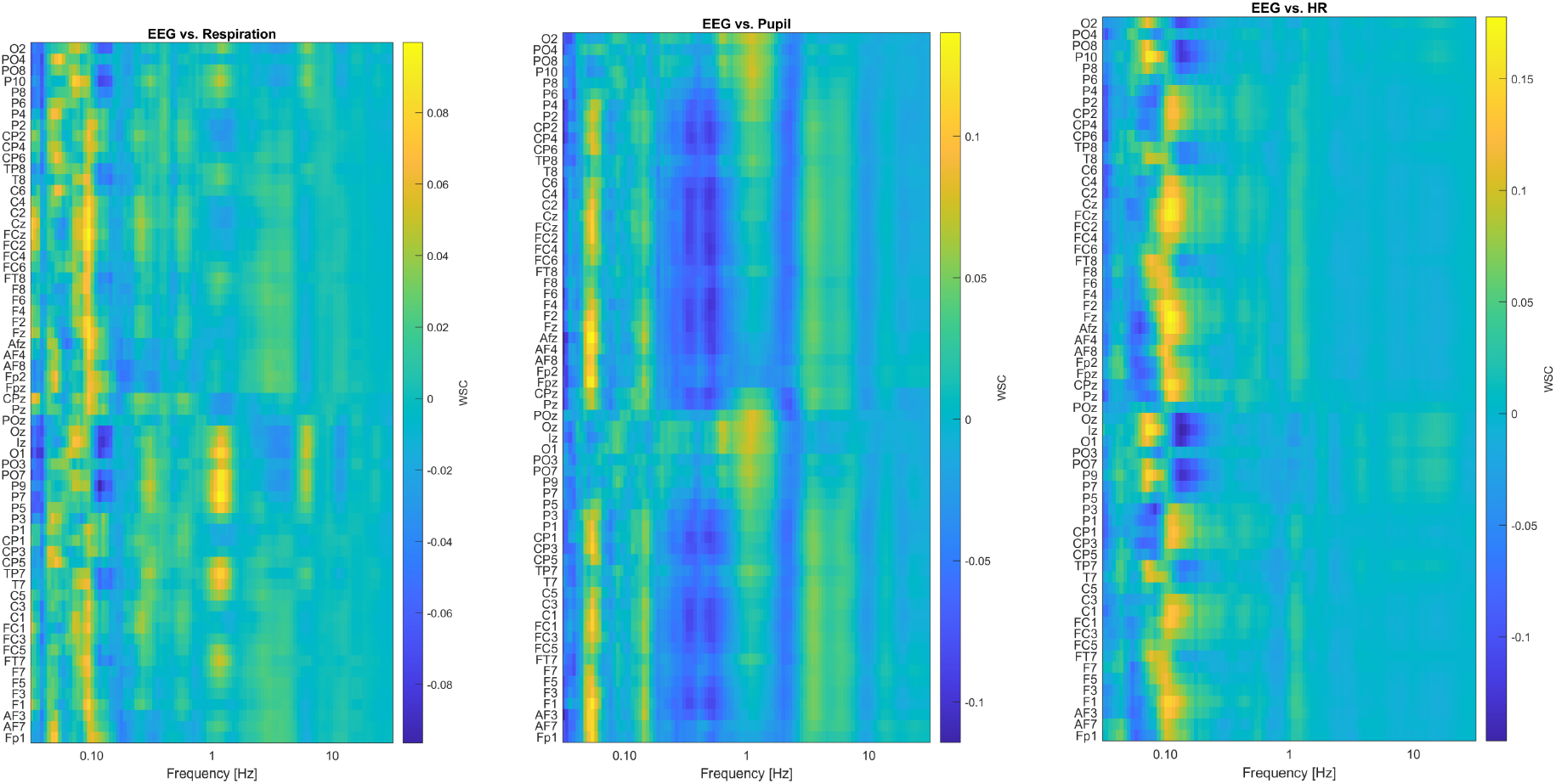
Spatial distribution across the scalp of WSC, i.e. correlation between raw EEG and respiration, pupil and HR signals, resolved by electrode and frequency band.

**Figure S4:**
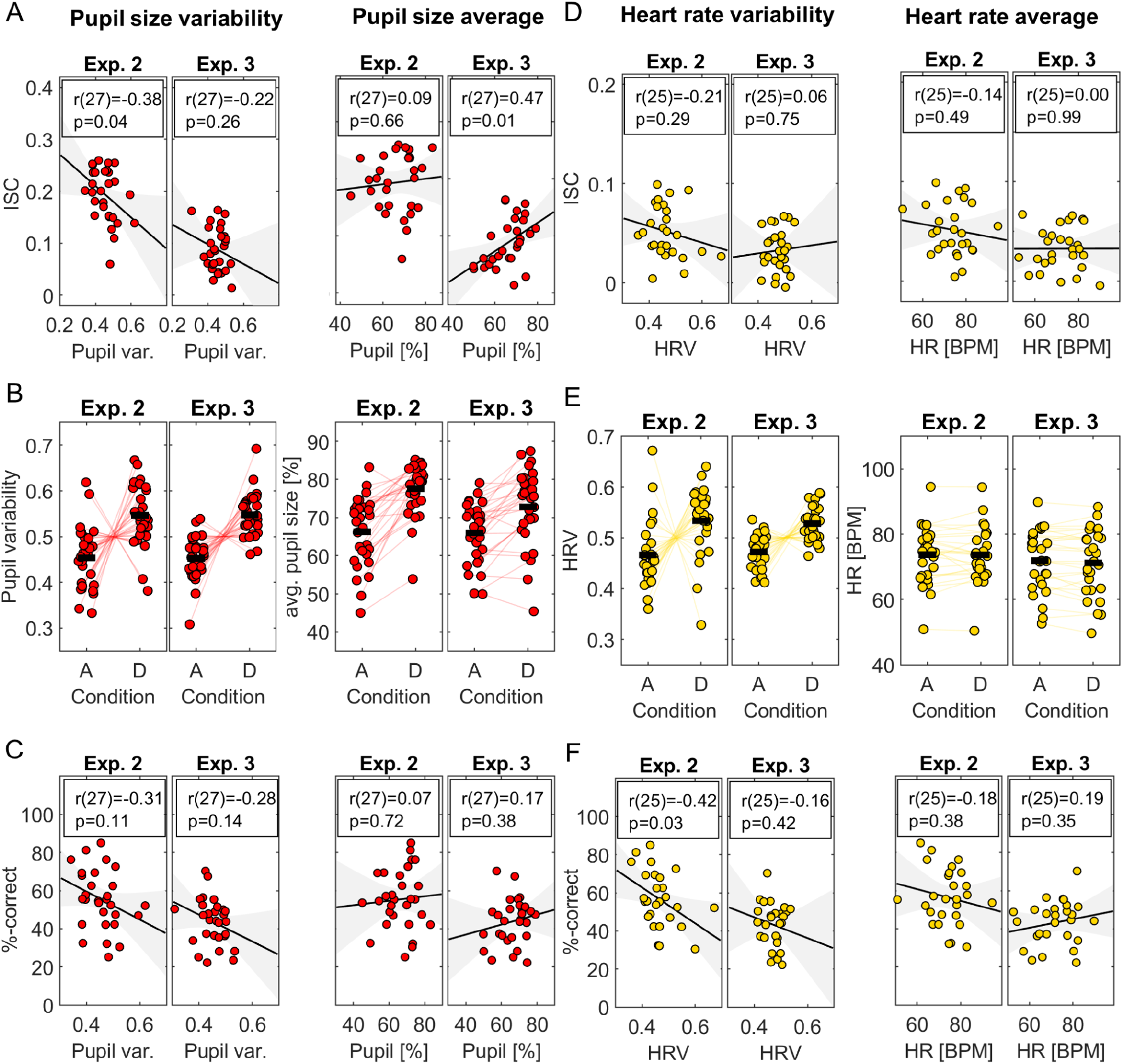
Relation between pupil size and heart rate with memory, ISC and attention: Pupil size, pupil variability, heart rate and heart rate variability were measured for each subject and video clip (time scale of minutes) and averaged over videos for Experiment 2 (Exp. 2) and Experiment 3 (Exp. 3) **A:** average pupil size and pupil variability for each subject compared to the ISC **B:** Pupil size and pupil variability in the attending and distracted conditions. Lines are connected between the average pupil size of a subject when they watched the 5 videos in an attending condition (A) and distracted conditions (D). **C:** relationship with memory across subjects for pupil size and pupil variability **D:** same as A) but now for heart rate and heart rate variability **E:** same as B) but now for heart rate and heart rate variability **F:** same as C) but now for heart rate and heart rate variability

**Figure S5:**
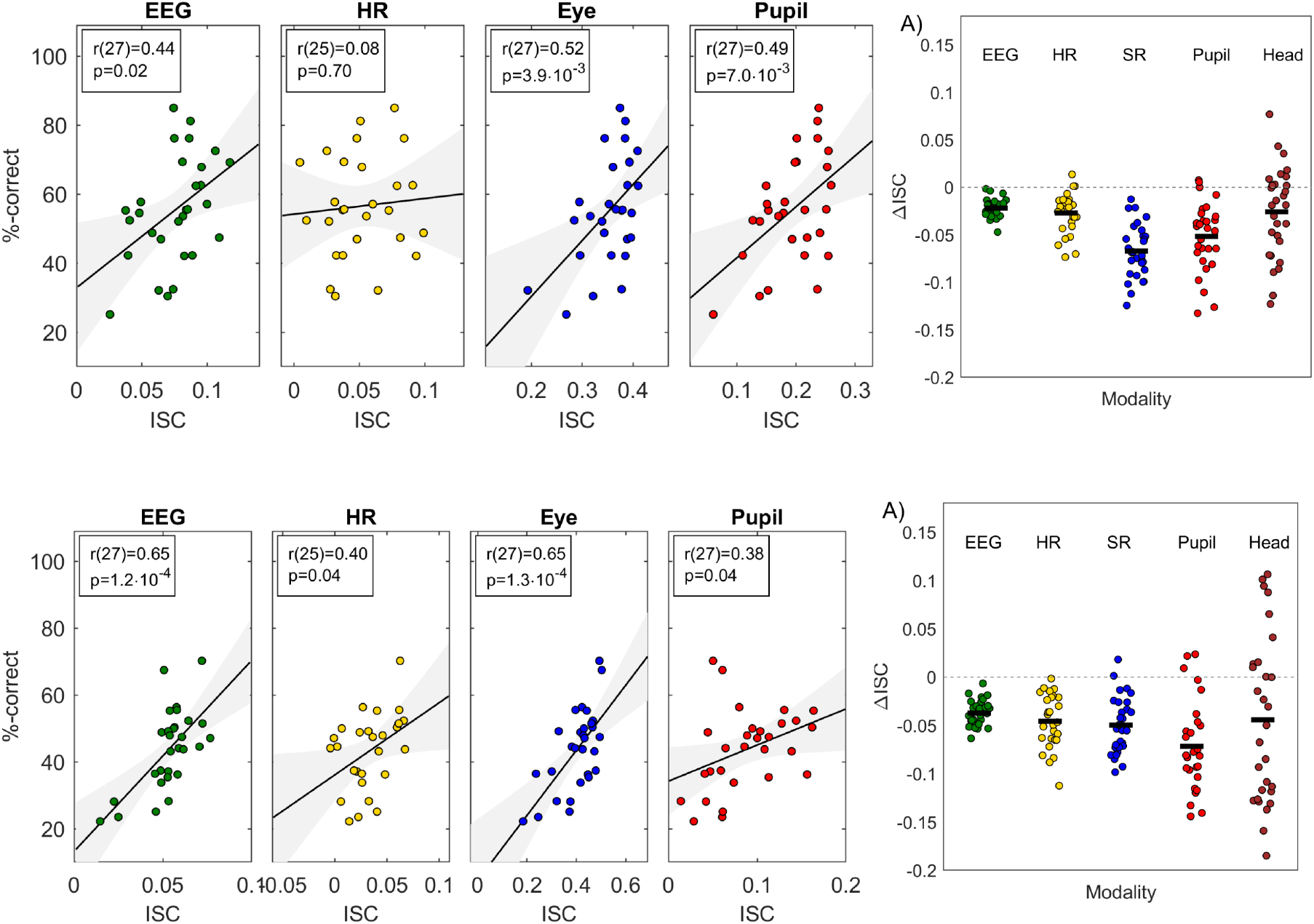
Correlation of ISC with memory and modulation with attention replicates over two additional experiments: First row: Data from Experiment 2 (N=29). For HR 2 subjects were removed due to bad signal quality. Second row: Data Experiment 3 dataset (N=29). For HR 2 subjects were removed due to bad signal quality. Each point is one subject. Right panels show change in ISC (D condition - A condition) for each subject.

**Figure S6.**
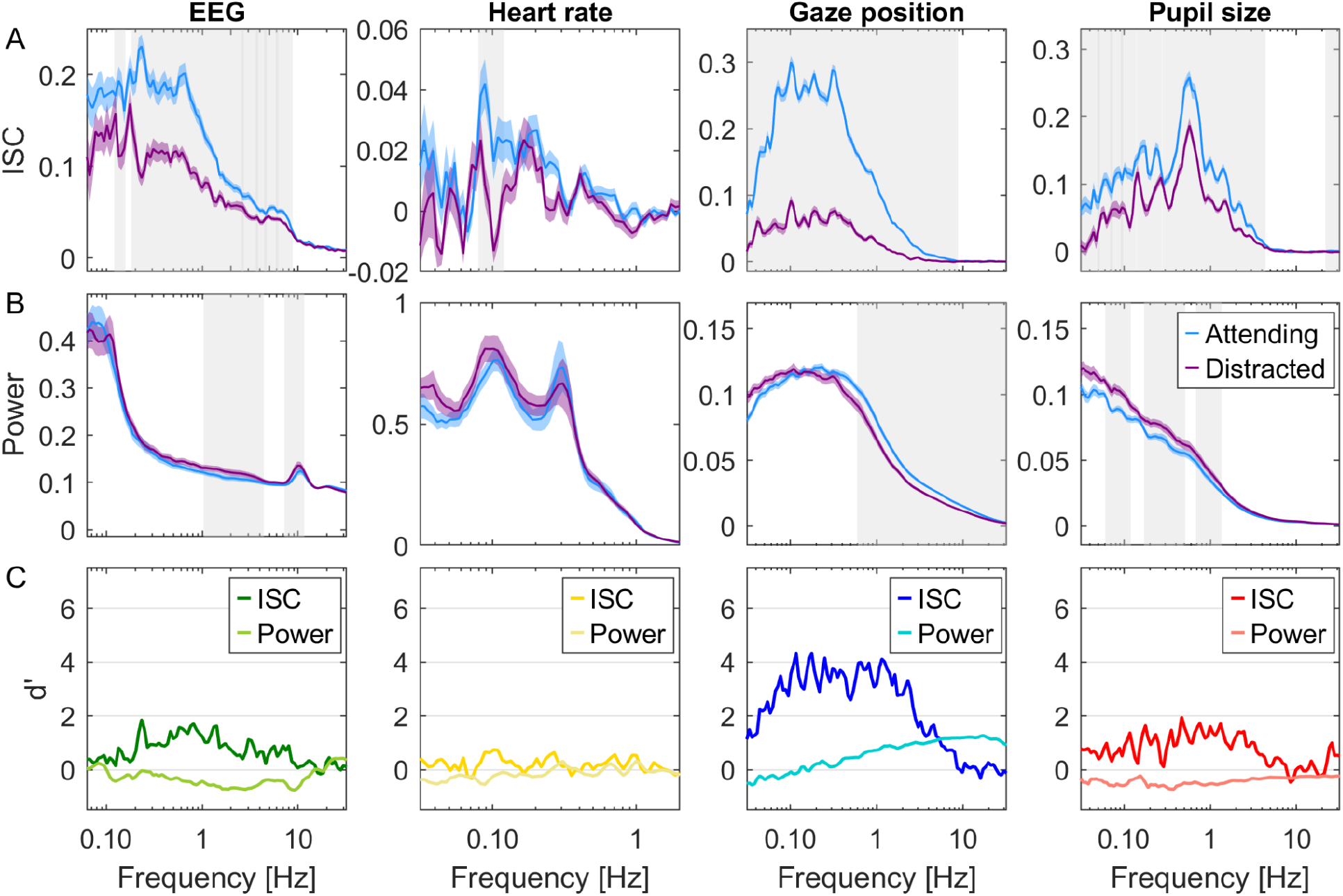
Attentional modulation of ISC and Power resolved by frequency. Same as Fig. 5, but for data from Experiment 3.

